# Deployable high-fidelity metagenome binning at scale with QuickBin

**DOI:** 10.64898/2026.01.08.698506

**Authors:** Brian Bushnell, Juan C. Villada

## Abstract

Reconstructing genomes from metagenomic assemblies is foundational to microbiome research, yet metagenome binning remains constrained by a persistent trade-off between genome fidelity and deployable throughput. Many high-accuracy approaches rely on GPU-intensive workflows, marker-gene-informed postprocessing, or resource demands that limit reproducible use at catalog scale. Here we present `QuickBin`, a CPU-native, marker-free binning algorithm designed to recover near-complete, ultra-low-contamination metagenome-assembled genomes (MAGs) under practical compute constraints. `QuickBin` couples a GC-coverage spatial index (BinMap) with a hierarchical, early-exit Oracle cascade of similarity tests (scalar composition/coverage filters followed by SIMD-accelerated k-mer comparisons), invoking a compact neural network only for the small fraction of ambiguous candidate merges. Across synthetic communities evaluated by both marker-based estimates and contig-origin ground truth, `QuickBin` yields high recovery while maximizing the fraction of sequence assigned to uncontaminated bins. In a benchmark of 297 diverse real metagenomes, `QuickBin` completed all runs and recovered more MAGs meeting ≥95% completeness and ≤1% contamination than widely used CPU binners and compute-intensive alternatives that frequently failed to finish under standardized limits. `QuickBin` provides a practical path to reproducible, high-fidelity genome-resolved metagenomics at scale for downstream comparative analyses and data products that depend on low-artifact genome reconstructions. The full software package is open-source and available for download at https://bbmap.org, `GitHub` repo (https://github.com/bbushnell/BBTools), or `Docker` container (https://hub.docker.com/r/bryce911/bbtools).

## 2. Introduction

Genome reconstruction from metagenomic data has defined our modern understanding of microbial evolution, metabolism, and ecology by enabling analysis of uncultivated lineages that are present in many environments (Bowers et al. 2017; Hug et al. 2016; Rinke et al. 2013; Nayfach et al. 2021; Sun et al. 2025). Metagenome-assembled genomes (MAGs) are widely used in microbiome research to infer functional potential across diverse biomes, from the human gut (Almeida et al. 2021; Pasolli et al. 2019) to deep-sea sediments (Nayfach et al. 2021) and soil ecosystems (Bahram et al. 2018). MAGs now support comparative microbiome analyses, ecological inference, and the construction of large genome catalogues, ruled by community quality standards such as the Minimum Information about a Metagenome-Assembled Genome (MIMAG) (Bowers et al. 2017) that formalize expectations for completeness and contamination that enable robust downstream analyses.

A central step in this workflow is metagenomic binning, a process that consists in the assignment of assembled contigs into genome bins using signals such as sequence composition, differential coverage across samples, and assembly-graph connectivity (Sedlar et al. 2017). Over the past decade, a succession of algorithms has improved the resolution of metagenomic binning. Early composition-based methods (Dick et al. 2009) were succeeded by differential coverage approaches like `MetaBAT2` (Kang et al. 2019) and `MaxBin2` (Wu et al. 2016), which remain widely used standards. More recently, the integration of deep learning and variational autoencoders, exemplified by `VAMB` (Nissen et al. 2021), `SemiBin2` (Pan et al. 2022), and `ComeBin` (Wang et al. 2024), has further refined binning accuracy by leveraging complex feature embeddings. These advances have facilitated the generation of massive genomic resources, including the Unified Human Gastrointestinal Genome (UHGG) catalog (Almeida et al. 2021), the Genomes from Earth’s Microbiomes (GEM) catalog (Nayfach et al. 2021), the global species-level repositories (Pasolli et al. 2019; Almeida et al. 2019); with newer bigger catalogs and repositories for MAGs like MGnify (Gurbich et al. 2023), gcMeta 2025 (Sun et al. 2025), and SPIRE (Schmidt et al. 2023) collectively housing millions of MAGs.

However, the rapid expansion of sequencing throughput has introduced significant computational and quality control challenges. Current binning strategies frequently face a trade-off between scalability and precision (Meyer et al. 2022). As datasets approach even larger scales, the computational cost of all-versus-all alignment and high-dimensional clustering becomes a bottleneck for existing tools (Wang et al. 2023; Nissen et al. 2021). Furthermore, recent quality evaluations of MAGs have revealed that a substantial fraction of publicly available MAGs contain undetected contamination, particularly when automated pipelines prioritize recall over precision (Shaiber and Eren 2019; Han et al. 2025).

Recent foundation-model studies indicate that large-scale pretraining on nucleotide sequences is feasible and can produce representations that transfer to downstream genomic tasks (Dalla-Torre et al. 2025; Nguyen et al. 2023, 2024), while related protein-language-model studies have shown how model behavior depends on the statistical structure of the training data (Rives et al. 2021; Lin et al. 2023). As such efforts expand, the community benefit of genome resources is no longer limited to comparative genomics and ecology. It also extends to enabling reliable genomic AI, where minimizing chimeric artifacts and contamination in training genomes is a reasonable prerequisite for model robustness (Whang et al. 2023; Bileschi et al. 2022; Jain et al. 2022). The imperative for high-fidelity genomic data has intensified with the advent of artificial intelligence in biology. Genomic foundation models and large language models (LLMs) trained on biological sequences require data of exceptional purity to learn accurate representations of syntax and function (Nguyen et al. 2023; Ji et al. 2021). Contaminated, fragmented, or marker-gene biased construction of MAGs risk introducing noise that can propagate hallucinations in generative models, or obscure subtle evolutionary signals (Whang et al. 2023; Yan et al. 2024; Bileschi et al. 2022).

Existing MAG catalogues provide broad coverage, but MAG contamination levels are variable. Growing reliance on MAGs as reusable references elevates the importance of bin purity beyond what is captured by conventional “high-quality” thresholds as MIMAG high-quality genomes allow up to 5% contamination. This motivates development of new tools that aim for high completeness and, especially, low contamination to support use cases that require reliable training data (Lamurias et al. 2022; Uritskiy et al. 2018; Sieber et al. 2018).

Here, we present `QuickBin`, a resource-efficient metagenome binning algorithm designed to address scalability and genome-fidelity challenges in large-scale genome-resolved metagenomics. `QuickBin` leverages a small, pretrained feed-forward neural network, only as a final adjudicator for plausible pairs prioritized by computationally inexpensive heuristics. This enables rapid binning on commodity CPU-only nodes or laptops, on any operating system or architecture, with no compilation or installation dependencies other than Java (or Docker), while prioritizing recovery of near-complete, contamination-free genomes. `QuickBin` reduces the effective computational burden by combining two ideas. First, it uses BinMap, a GC-coverage spatial index that restricts comparisons to local neighborhoods in feature space rather than performing global all-versus-all evaluations. Second, it uses a hierarchical Oracle decision cascade that aggressively filters candidate merges using inexpensive composition and coverage checks before applying more sensitive SIMD-accelerated k-mer similarity tests; a compact feed-forward neural network is used only as a final adjudicator for the small subset of ambiguous candidate pairs that survive earlier filters. This architecture enables `QuickBin` to operate robustly on commodity CPU infrastructure with low memory budgets while remaining marker-free, preserving broad applicability across taxa (including Eukaryotes).

We benchmark `QuickBin` against against widely used and recent state-of-the-art binners spanning coverage-and-composition binners (for example `MetaBAT2`) (Kang et al. 2019), a VAE-based binner (`VAMB`) (Nissen et al. 2021), self-supervised and contrastive-learning binner (`SemiBin2`) (Pan et al. 2023), and a contrastive multi-view representation learning binner (`COMEBin`) (Wang et al. 2024); and we evaluate with both `CheckM2` (Chklovski et al. 2023) tiers and ground-truth mapping. Across synthetic benchmarks evaluated by `CheckM2` and by ground-truth contig-origin mapping, `QuickBin` consistently achieves strong genome recovery together with high bin purity, and in large multi-coverage settings it attains near-complete recovery with near-complete clean-bin signal. In real metagenome benchmarks spanning diverse biomes, `QuickBin` supports robust recovery of high-fidelity MAGs, including strong performance in the stringent regime of completeness ≥95% and contamination ≤1%, while maintaining orders of magnitude lower runtime and memory requirements than other binners. Our results show that `QuickBin` is a practical tool for scalable generation of high-confidence MAG sets for comparative microbiome analyses and for downstream applications that depend on low-artifact genome reconstructions, including genomic machine learning.

## 3. Results

`QuickBin` was developed to enable high-fidelity genome recovery in large-scale metagenomic collections where algorithmic accuracy is often compromised by operational constraints, and to accurately deconvolve high-quality genomes from non-axenic or contaminated isolates and low-complexity commensal libraries. Standard workflows frequently sacrifice bin purity or completeness when assemblies are massive or compute resources are limited. `QuickBin` addresses these challenges by integrating GC-content, multi-sample coverage, and graph connectivity signals to prioritize conservative contig merging (Figure 1a). This conservative approach is a prerequisite for recovering genomes that meet beyond MIMAG standards (≤1% contamination), ensuring they are suitable for high-confidence interpretation and the training of genomic foundation models.

**Figure 1.**
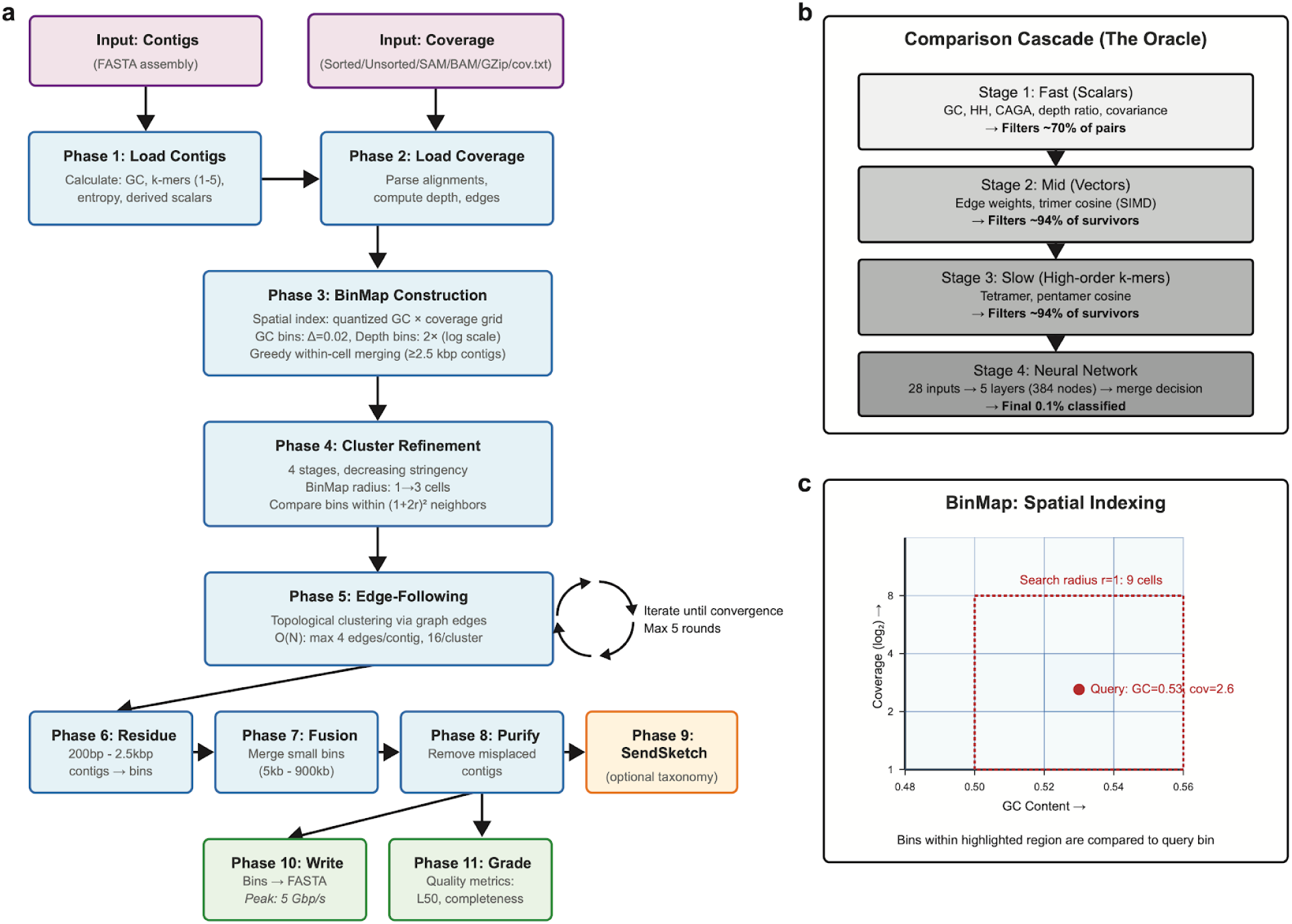
QuickBin algorithmic workflow and comparison methodology. **(a)** The main algorithm flow through 11 phases. Purple boxes indicate input data; blue boxes represent processing phases; green boxes show output generation. Cycle arrows indicate iterative refinement loops. Phases 1-2 load and preprocess input data, computing metrics concurrently via multithreaded pipelines. Phase 3 constructs the BinMap spatial index (see Figure 1c for details). Phases 4-5 perform primary clustering via spatial (BinMap) and topological (graph edge) similarity, iterating until convergence. Phases 6-8 process small contigs, merge fragmented genomes, and correct greedy algorithm errors. Phases 9-11 handle optional taxonomic annotation, file output, and quality assessment. **(b) The Oracle comparison cascade.** Bin pairs are filtered through four stages of increasing computational cost: (1) scalar metrics eliminate ∼70% of pairs; (2) trimer frequency vectors (SIMD-accelerated) eliminate ∼94% of survivors; (3) tetramer/pentamer frequencies eliminate another ∼94%; (4) the remaining 0.1% are classified by a neural network trained on synthetic metagenomes. This cascade reduces effective complexity from O(N^2^) to near-linear. **(c) BinMap spatial indexing structure.** The multidimensional index quantizes GC content (Δ=0.02) and coverage depth (2× logarithmic bins) to partition feature space. The example query bin (red dot; GC=0.51, coverage=1.3) is compared only to bins in neighboring cells within search radius r. For r=1, this encompasses (1+2r)^2^=9 cells (red dashed box), dramatically reducing the number of comparisons from all-pairs O(N^2^) to spatially localized subsets. **\* Abbreviations:** GC, guanine-cytosine content; HH, homo/heteropolymer ratio; CAGA, inter-GC transition affinity; SIMD, single instruction multiple data; HQ, high-quality; ANI, average nucleotide identity; SSU, small subunit ribosomal RNA; O(N), computational complexity order notation.

The framework’s discriminatory power is concentrated in the Oracle cascade (Figure 1b), a hierarchical decision pipeline designed to prevent the “hallucinations” of chimeric artifacts. To maintain fidelity without the computational exhaustion of all-versus-all deep learning, the Oracle triages comparisons through a series of early-exit heuristics. It first rejects candidates with divergent GC content, depth ratios, or depth covariance. Only those surviving these initial filters are subjected to hierarchical k-mer cosine similarity tests (trimers, tetramers, and pentamers). The pretrained feed-forward neural network acts as the final adjudicator, activated only for the small subset of plausible, ambiguous contig pairs that pass all prior heuristic thresholds.

To achieve scalability, `QuickBin` utilizes BinMap spatial indexing (Figure 1c). By quantizing GC and multi-sample coverage into a multi-dimensional grid, BinMap constrains candidate comparisons to local neighborhoods. This avoids the quadratic complexity of global comparisons, allowing the algorithm to enforce stringent merge criteria across datasets with millions of contigs while remaining within limited memory and walltime bounds.

### 3.1. QuickBin Algorithm

`QuickBin` solves the computational exhaustion bottleneck in metagenomic binning by prioritizing evidence-based triage. Unlike deep-learning binners that subject every contig pair to expensive neural-network evaluations, `QuickBin` utilizes a hierarchical Oracle cascade.

This multi-phased approach evaluates sequence composition, coverage, and graph connectivity through a series of early-exit filters. Computationally inexpensive scalars, such as GC content and dimer-derived metrics (HH, CAGA), are employed first to immediately discard dissimilar pairs. This strategy reserves more sensitive and expensive comparisons, such as trimer frequency cosine difference and the terminal neural network layer, for only the most ambiguous surviving pairs. By combining this hierarchical triage with BinMap spatial indexing, `QuickBin` achieves a 14-second runtime and 98.8% clean-bin purity on a single CPU, providing a high-fidelity alternative to resource-intensive black box models.

#### 3.1.1. Calculated metrics

##### 3.1.1.1. Coverage

Read coverage depth for each contig is calculated from SAM or BAM files, which are loaded via multithreaded streaming then discarded, ensuring memory usage remains independent of the size or number of input alignment files. Coverage can be emitted as a compact text file (`cov.txt`) to allow rerunning without reprocessing SAM/BAM files. Coverage can be input to `QuickBin` as SAM or BAM, as sorted or unsorted, and the files can be uncompressed or `gzip`/`bgzip-`compressed. Coverage is calculated as:

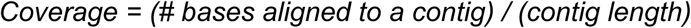

##### 3.1.1.2. Coverage Covariance

When multiple SAM/BAM files are provided, coverage for each is treated as an independent sample (e.g., a time or depth series) and tracked individually. Coverage cutoffs are then applied on a per-sample basis such that if contigs A and B have similar depth in sample 1 but dissimilar depth in sample 2, they will not be clustered together. Because greater numbers of samples allow greater specificity from coverage covariance alone, with increasing numbers of samples, other cutoffs such as GC difference are relaxed.

To ensure the accuracy of this technique, an entropy calculation determines the number of samples-equivalent of information content. For example, the same sample sequenced across two sequencing lanes and presented as 2 SAM files would have their coverage tracked independently but would internally be considered as approximately 1.05 samples of information content due to minor discrepancies in coverage, since they would covary almost perfectly.

##### 3.1.1.3. GC Content

Calculated as:

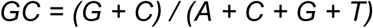

This is used extensively as an efficient scalar threshold when comparing contigs. GC content is calculated simultaneously with all other short k-mer frequencies during the multithreaded assembly loading phase. As GC content stabilizes toward the genome average with increasing contig lengths, the allowable difference between two contigs is determined by their length, as with other metrics.

##### 3.1.1.4. Short k-mer Frequency

k-mer counts of length 1-5 are calculated during contig loading and stored in `int[]` arrays associated with each contig, as raw counts for k=1-2 and canonical counts for k=3-5 (k=5 is only calculated for contigs ≥2 kbp). The actual comparisons use a pair of scalars derived from the dimers; SIMD-accelerated cosine difference (*1 − cosine similarity*) for trimers, tetramers, and pentamers; and additionally, GC-compensated cosine difference for tetramers strictly as an input to the neural network. These are stored as counts instead of frequencies to allow fast and exact merging of contigs and clusters with updated count arrays.

The GC compensation calculates frequencies of k-mers relative to only the subset of k-mers with the same GC content; for example, the GC-compensated frequency of AAAA would be calculated as:

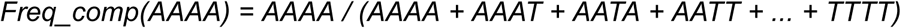

(using only k-mers where GC=0)

##### 3.1.1.5. Entropy

GC-compensated entropy is calculated while loading contigs, by calculating tetramer frequencies over a sliding window of 150 bp and averaging the window entropy over the full contig. The GC compensation was developed using empirical observation of simulated random sequences at varying GC content, such that the range is 0-1 across the GC spectrum rather than extreme GC sequence having a lower maximum entropy and neutral-GC sequence having higher maximum entropy.

##### 3.1.1.6. Derived Scalars

Strandedness, HH, and CAGA are all derived from dimer frequencies:

- Strandedness: Ratio of sum of minority dimers to the sum of their complements. For example, if |AC| > |TG| and |GC| > |CG|, then AC and GC would be in the denominator, and TG and CG in the numerator; a value near 1 indicates a lack of dimer frequency strand preference (as expected in random sequence), while a lower value indicates bias. This metric is inherently GC-independent.
- HH (Homo/Het Polymer Ratio): Calculated as: HH = (AA + CC + GG + TT) / (AA + TT + AT + TA + CC + GG + CG + GC) A value over 0.5 indicates that homodimers are more common than heterodimers, amongst dimers with GC=1 or GC=0. This metric is inherently GC-independent.
- CAGA (Inter-GC Transition Affinity): Calculated as: CAGA = 0.5 × (1 + (CA + TG − GA − TC) / (AC + AG + CA + GA + TC + TG + CT + GT)) Values above 0.5 show a bias toward CA and TG, while values below 0.5 show a bias toward GA and TC. The numerator terms and signs were selected using statistical analysis to determine which dimer frequencies were most strongly correlated or anticorrelated, along with which had the greatest interspecies variance and lowest intraspecies variance. This metric is inherently GC-independent.

These three scalars form GC-independent stable dimensions which are inherent properties of a contig alone, and thus can be calculated independently of coverage or other contigs—unlike GC/Coverage plots or tetramer frequency PCA plots, which rely on external information for dimensions. Of the derived scalars, HH and CAGA differences are used as cutoffs to shortcut more expensive comparisons, while strandedness is only used as a neural network input.

##### 3.1.1.7. Edges

Assembly graph connectivity is calculated using paired read mapping, in cases where each read maps to a different contig. These edges are weighted based on the number of spanning pairs compared to the depth of the contig, and depending on how many outward edges a contig has per end. The presence of edges between contigs does not force them into the same cluster, but is used to lower other thresholds and recruit small contigs that would not otherwise be compared at all.

#### 3.1.2. Algorithm Phases

##### 3.1.2.1. Phase 1: Load Contigs

`QuickBin` starts by loading all contigs with a multithreaded file reader; as contigs are read from disk, additional threads competitively calculate contig-specific metrics such as GC, entropy, and k-mer counts. This architecture achieves 590 Mbp/s throughput at an observed 3,100% CPU utilization on a 32-core system, loading a 4 Gbp assembly in 6.7 seconds. Optionally, 16S/18S SSUs are gene-called concurrently. K-mer counting uses an optimized function that counts 1, 2, 3, 4, and 5-mers simultaneously.

##### 3.1.2.2. Phase 2: Load Coverage

`QuickBin` can be run with or without coverage information. If no alignment information (SAM, BAM, or cov files) is provided, coverage estimates from the assembler will be automatically parsed from contig headers (supporting `Tadpole` [https://bbmap.org/tools/tadpole]), `SPAdes` (Nurk et al. 2017), and `MetaHipMer` (Hofmeyr et al. 2020) headers). In the case of nonuniform amplification such as MDA, coverage is misleading and the “flat” flag should be used to ignore contig depths, while graph connectivity can still be used based on paired alignment results. However, binning quality will suffer in the absence of coverage information so it should always be used when available and accurate.

When loading SAM or BAM files, multiple files (up to the number of threads) are loaded simultaneously. Each file uses multithreaded `bgzip` decompression and multithreaded file parsing. As a result, bgzipped files (the output of all BBTools when a `.gz` extension is used) are loaded faster than raw uncompressed SAM due to filesystem bandwidth constraints, and even a single BAM file can saturate 13 cores, measured at 1,420 Mbp/s. Nevertheless, the coverage and graph connectivity can be stored in a small text file with the “`covout`” flag, which loads almost instantly when reprocessing the same assembly with different parameters. Without a pregenerated coverage text file, coverage loading is usually the longest phase.

##### 3.1.2.3. Phase 3: BinMap Creation

`QuickBin` represents sequences as Bin objects, with subclasses Contig or Cluster (a mutable set of multiple Contigs). Each Bin has fields such as GC, HH, trimers, and a depth array (per sample) which are compared between bin pairs. During clustering, Contigs or Clusters can be merged to form larger Clusters, causing the values of the new member to be added to the fields of the new or modified Cluster. For example, merging two Contigs would sum the dimer count arrays, perform a size-weighted average of the depth arrays, and force recalculation of GC from monomer counts. Thus, to allow multithreading and prevent problems of synchronization on changing global state, most clustering phases are performed in two subphases: multithreaded comparison of specific Bin pairs, followed by recursive merging of all pairs identified as sufficiently similar. Because this process creates new or modified Bins, the process is typically repeated for a fixed number of rounds or until convergence. The initial creation of the data structure used for most of these phases, though, is singlethreaded, using only a single round and combining comparison and merging greedily.

The BinMap itself is a conceptual GC/cov grid (or with multiple samples, a GC/cov1/cov2 cube) containing a set of lists corresponding to quantized GC or coverage levels. GC is quantized linearly in increments of 0.02 and depth is quantized exponentially by a factor of 2. Thus, a Bin with GC=0.501, cov=3.5 would be in the same list as another with GC=0.505, cov=3.7; adjacent to a Bin with GC=0.523, cov=3.3; and 2 coverage steps plus 1 GC step away from a Bin with GC=0.492, cov=9.9.

When creating the BinMap from the input assembly, Contigs are processed in size-descending order and only compared within their own cell. Thus, the first Contig in a cell will be compared to nothing and simply added to the list. Subsequent Contigs will be compared to each existing Bin in the list, and merged with the first one passing the comparison cutoff, or else appended.

Therefore, this phase is O(N^2^) and would require N(N-1)/2 comparisons if all Contigs were added, no merges were performed, and all Contigs were in the same GC/cov cell. However, in practice it is very fast because elements in the same cell often merge, the metagenomic feature space is well divided by these metrics, and only Bins of at least 2.5 kbp are added to the BinMap.

##### 3.1.2.4. Phase 4: Cluster Refinement

The primary clustering occurs here, with 4 stages of gradually decreasing stringency. Every Bin is compared to all other eligible Bins. Eligibility is decided using the BinMap with a maximal cell distance, which is the number of cells away the two are in the quantized GC or coverage dimension. This range varies from 1 to 3. With range 1, a Bin would be compared to Bins in up to 9 lists: its own list, and all lists with ±0.02 GC or ±1 depth (on a log_2_ scale). The number of lists used for comparison is up to (1+2*range)^2^, so range 3 uses up to 49 lists of Bins. The comparison is multithreaded, followed by single-threaded merging of identified good pairs to prevent concurrent modification.

##### 3.1.2.5. Phase 5: Edge-following

If mapped paired reads were provided, or embedded in the cov file, edge-following is performed; all contig pairs with mapped spanning read pairs of sufficient depth are compared to each other, and sufficiently similar pairs are merged into the same cluster. The comparison methodology between pairs is the same in all clustering phases, so the difference between this clustering phase and refinement is simply how the pairs being compared are selected.

For QuickBin, an all-to-all comparison would be close to twice the number of comparisons for all pairs (N(N-1)/2) because it compares not only contig pairs but cluster pairs as they grow.

However, this is still O(N^2^) rather than O((2^N^)^2^) (all subsets versus all subsets) since as a mostly greedy algorithm (adding a contig to a cluster is generally irreversible), each new cluster needs to be compared to at most O(N) elements, and clusters can only be formed or changed O(N) times. The edge-following phase, however, is only O(N) expected, because each Bin compares to at most a constant number of others: 4 for Contigs and 16 for Clusters, using their strongest edges. As with refinement, pairs connected by edges are compared multithreaded, then merges are made singlethreaded. This repeats until convergence or 5 rounds have transpired.

##### 3.1.2.6. Phase 6: Residue Processing

While the initial BinMap construction and refinement was restricted to contigs of at least 2.5 kbp, this phase uses the “residue“: all contigs above the “mincontig” cutoff (default 200 bp) but smaller than the BinMap threshold of 2.5 kbp. Each residual contig is compared only to Bins already in the map, not to each other, and added to the best-matching bin according to the Oracle (if it passes the cutoffs). However, since stringency is higher for shorter contigs, most are not added.

##### 3.1.2.7. Phase 7: Fusion

In an attempt to combine genomes split over a few large bins, fusion compares bins in the size range of 5 kbp-900 kbp (below the expected size of a complete bacterial genome) to larger bins. The algorithm is otherwise the same as refinement apart from lower stringency and size thresholds.

##### 3.1.2.8. Phase 8: Purification

This is the only subtractive phase, attempting to correct errors made by the greedy algorithm by removing Contigs. Each Contig is removed from its Cluster and then compared to what remains (which may have changed since it was initially added). If the Oracle decides that it no longer belongs, it is removed. This is also multithreaded; each Cluster is completely handled by a single thread since they are being modified.

##### 3.1.2.9. Phase 9: SendSketch

Optionally, output Bins are sketched using BBSketch (https://bbmap.org/tools/sendsketch.html) and the sketch sent to the Sketch server at JGI (https://bbmap.org/services/bbsketch), where it is compared to all RefSeq (Goldfarb et al. 2025) organisms for taxonomic assignment. This information does not affect binning, but the predicted taxonomy of each bin will be printed in the report file for convenience.

##### 3.1.2.10. Phase 10: Writing

Bins are now written to disk with a multithreaded writer as a single FASTA file each, using one thread per file. To prevent thrashing on low-performance filesystems the default number of concurrent files is 1, but throughput increases by 25% when set to 2 on a high-performance flash SSD and over 3× faster when writing to RAM disk (/dev/shm/) with 4 threads, peaking at 5 Gbp/s. If the output files are given a `.gz` extension they will be bgzipped with multiple threads per file. Bins below mincluster (default 50 kbp) are written to a single chaff FASTA file if the chaff flag is set (it defaults to false).

##### 3.1.2.11. Phase 11: Grading

The completed bins are graded using `GradeBins` (https://bbmap.org/tools/gradebins). Synthetic data with TaxID labels will have completeness and contamination determined for all bins for the report, and a summary such as number of HQ bins and total score will be printed to the screen. For unlabeled data, only statistics such as L50 (size of the 50th percentile bin) and percent sequence recovery are reported.

#### 3.1.3. Comparison methodology

The actual comparison of Bins is the same in all clustering phases, though the stringency varies (generally becoming less stringent in later phases to compensate for the algorithm being greedy). Thus, the main difference between phases is determining which pairs to compare. Comparisons are performed by an Oracle object, which, subject to the desired stringency, makes the merge-or-not decision for a pair of Bins.

First, the GC, HH, and CAGA differences are computed using scalar subtraction. The maximal depth ratio (greater depth/lesser depth) is calculated across all samples. For example, with a single sample, this would simply be *max(a.cov, b.cov)/min(a.cov, b.cov)*, while for 2 samples it would be:

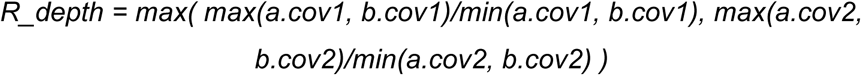

so it is selected from the sample in which their depth differs the most. To prevent extreme ratios in low-depth contigs a small constant is added to the numerator and denominator. If there are multiple samples, the coverage covariance is also calculated using the cosine difference of the normalized depth arrays. At this point, any of these metrics failing a cutoff will trigger an early exit from the comparison; this is called a “fast comparison” and typically eliminates approximately 70% of pairs from further consideration with just a few scalar operations. Note that most comparisons are never performed in the first place due to spatial indexing via the BinMap.

Second, graph edges between the pair are detected to adjust subsequent cutoffs; the cutoffs are less stringent for bins sharing edges. Some cutoffs are reevaluated in the presence of the additional connectivity information, and the cosine difference of the canonical trimer frequencies is calculated and subjected to its own cutoff. All cosine difference calculations are accelerated using 256-bit SIMD if available. This stage, “mid comparison”, eliminates approximately 94% of the remaining pairs.

Third, tetramer and (if both bins are over 40 kbp) pentamer frequencies are calculated and subjected to cutoffs. This stage, “slow comparison”, eliminates another 94% of the remaining pairs.

Finally, when the neural network is used, the surviving 0.1% of comparisons have the final decision made by the network. Otherwise the final decision is made using a composite of all metrics with a different cutoff. The network is a 75% connected dense randomly initialized feed-forward network with 28 inputs and 5 hidden layers, containing 384 nodes and approximately 25,000 weights, using mostly sigmoid and tanh activation functions with custom rslog (rotationally symmetric logarithm) as the final layer to better handle diminishing gradients. The network was trained using BBTools’ `train.sh` (https://bbmap.org/tools/train), using synthetic data from `Tadpole`-assembled TaxID-labeled contigs, coverage generated with `RandomReadsMG`, and `AllToAllVectorMaker` (`makequickbinvector.sh` [https://bbmap.org/tools/makequickbinvector]) which makes random subsets of contigs to compare for the purpose of making tsv files of vectors. The details of the input vector are listed in Table 1.

**Table 1.**
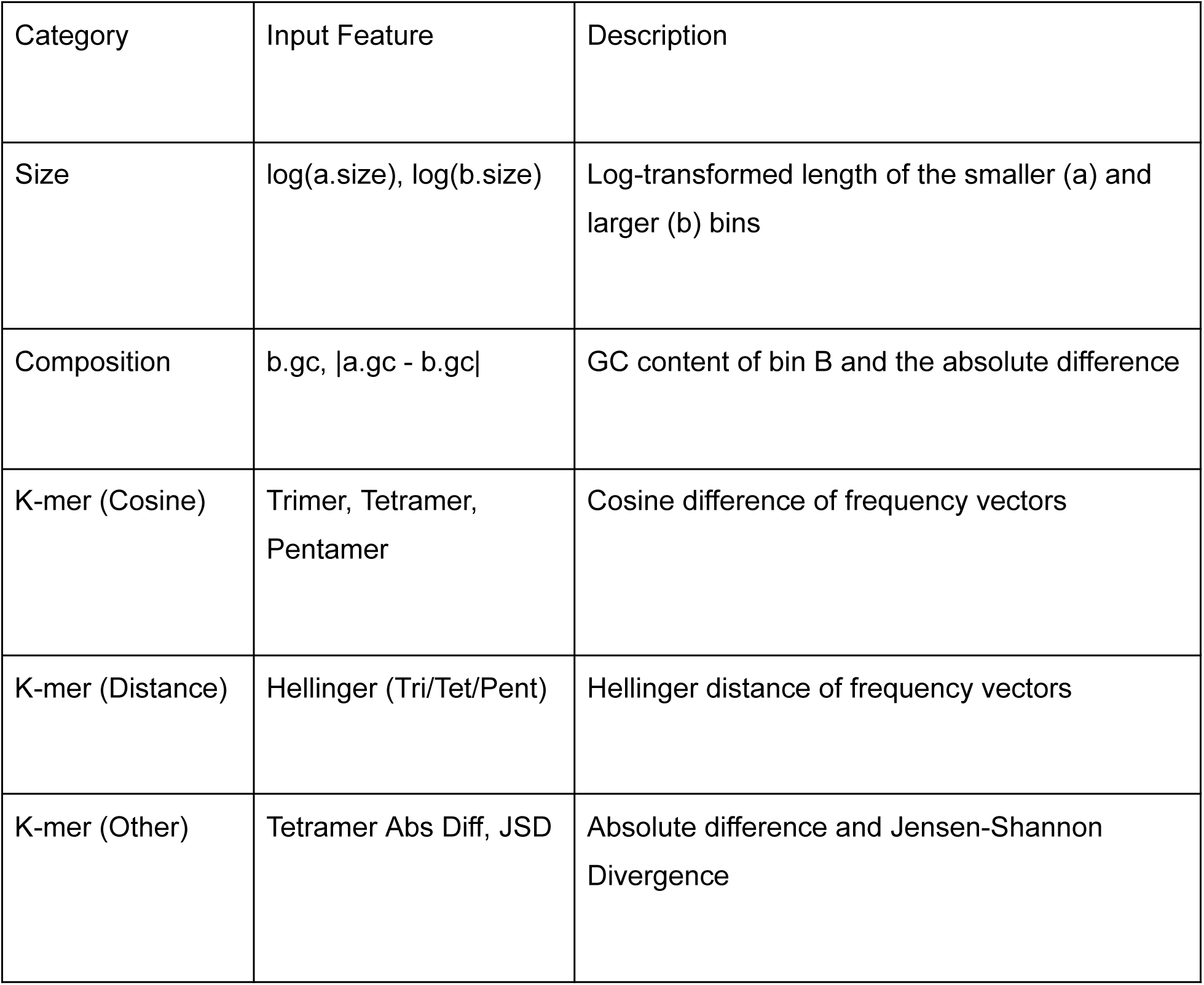
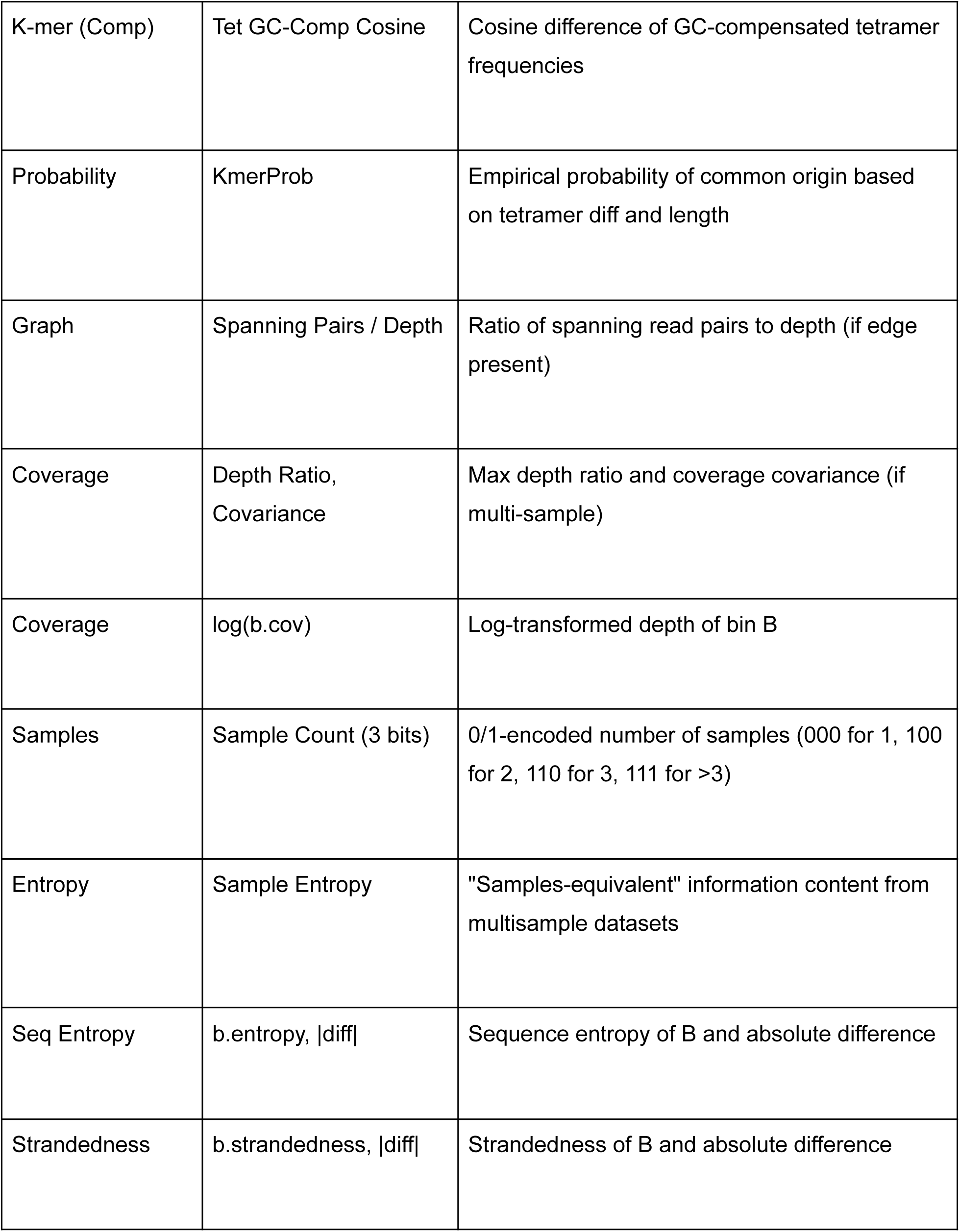
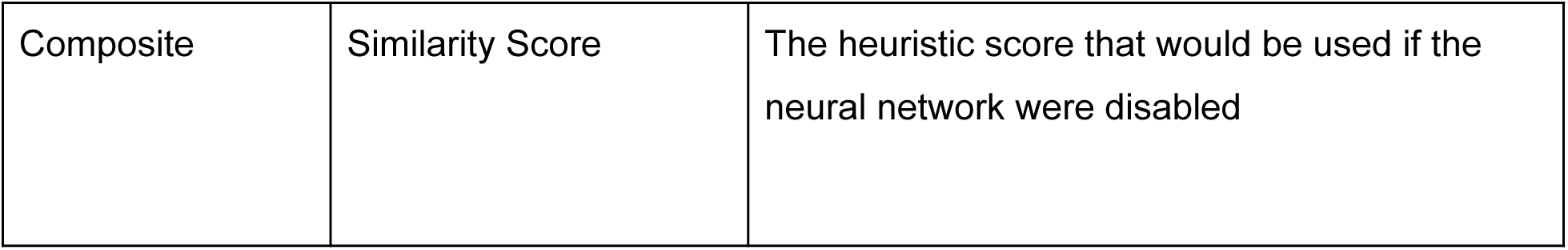
Neural network input vector for ground-truth assignment.

While `QuickBin` can be run without using the network, it dramatically improves the results on synthetic data (where the answer is known). See sections on “*Benchmarking `QuickBin` on synthetic data*” for more details.

In SSU mode (with the “ssu” flag), the final decision of merge/not merge can be vetoed when two bins passing all comparison cutoffs each have an identified SSU (typically 16S sequence, but 18S is also used). These sequences are aligned with the fast traceback-free QuantumAligner and the merge is rejected if the ANI is below 96%.

### 3.2. Selection of state-of-the-art binners for performance benchmarking

To establish a robust baseline for evaluating `QuickBin`, we first assessed the current landscape of metagenomic binning tools using performance metrics recently synthesized from a comprehensive benchmark by Han et al. (Han et al. 2025). An analysis of overall quality versus computational efficiency reveals a sharp trade-off between reconstruction quality and resource requirements of state-of-the-art algorithms (Supplementary Figure 1). Details on our ranking analysis based on the benchmarking done by Han et al. 2025 can be found in Supplementary Table S1-S3. Deep learning-based methods, specifically `COMEBin` (Wang et al. 2024) and `SemiBin2` (Pan et al. 2023), currently set the standard for reconstruction quality, ranking first and third, respectively. However, this precision comes at a substantial computational cost; `COMEBin` ranks lowest (13th) in both RAM and time efficiency, while `SemiBin2` similarly ranks in the bottom tier for computational speed. `MetaBinner` (Wang et al. 2023) was not considered as `COMEBin` was presented by the same group of authors as an improved tool.

**Supplementary Figure 1.**
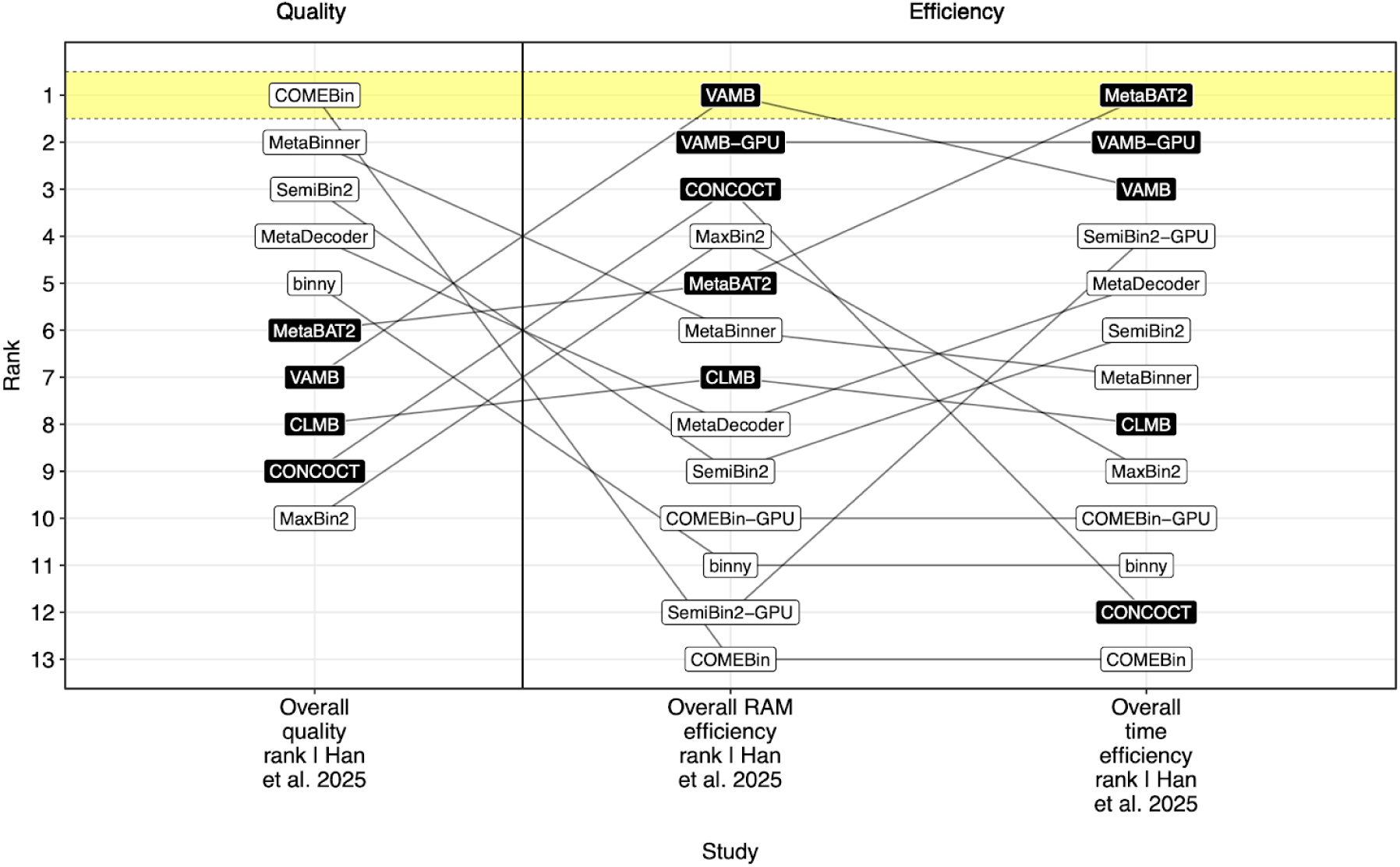
Comparative ranking of metagenomic binners across quality and efficiency metrics. Summary of performance rankings for state-of-the-art binning tools based on data from Han et al. (2025). Tools are ranked (1-13) across three distinct categories: Overall Quality, Overall RAM Efficiency, and Overall Time Efficiency. The chart highlights the performance trade-off between reconstruction quality and computational resource usage. `COMEBin` (highlighted in yellow, top left) achieves the highest quality rank (Rank 1) but the lowest efficiency ranks (Rank 13). Conversely, `MetaBAT2` (highlighted in yellow, top right) achieves the highest time efficiency (Rank 1) but places sixth in quality. `VAMB` (highlighted in yellow, top center) leads in RAM efficiency (Rank 1) while ranking seventh in quality. The connecting lines track the rank shifts of individual tools across metrics, illustrating the divergence between high-precision deep learning methods and high-efficiency clustering algorithms. Binners that use single-copy gene (SCGs) markers have a white background while binners that do not use SCGs have a black background.

Conversely, widely adopted tools such as `VAMB` (Nissen et al. 2021) and `MetaBAT2` (Kang et al. 2019) demonstrate superior resource efficiency. `MetaBAT2` ranks first in time efficiency and `VAMB` ranks first in RAM efficiency , yet they lag in reconstruction quality, ranking sixth and seventh, respectively. This trade-off, where high-fidelity reconstruction is inversely correlated with scalability, informed our selection of benchmarking targets. We therefore chose to compare `QuickBin` against `COMEBin` and `SemiBin2` to assess maximum achievable precision, and against `MetaBAT2` and `VAMB` to evaluate computational performance at scale.

### 3.3. Curation of a diverse global metagenomic dataset for real-world benchmarking

In order to assess the performance of `QuickBin` on real data, we collected a representative dataset of publicly available, unrestricted metagenomes sequenced by the Joint Genome Institute (JGI) and hosted on the IMG/M platform (Chen et al. 2023). This selection was designed to benchmark computational efficiency and binning fidelity across a wide spectrum of ecological and technical variability.

Geographically, the samples are distributed across four primary biomes: Aquatic, Engineered, Plant-associated, and Terrestrial (Supplementary Figure 2a).To ensure our benchmarking captured the challenges of modern sequencing workflows, we included data generated from both short-read (Illumina) and long-read (PacBio) platforms. The dataset spans a broad range of specific environmental subtypes, including deep subsurface aquifers, marine and freshwater systems, anaerobic bioreactors, and diverse soil horizons (Supplementary Figure 2b).

**Supplementary Figure 2.**
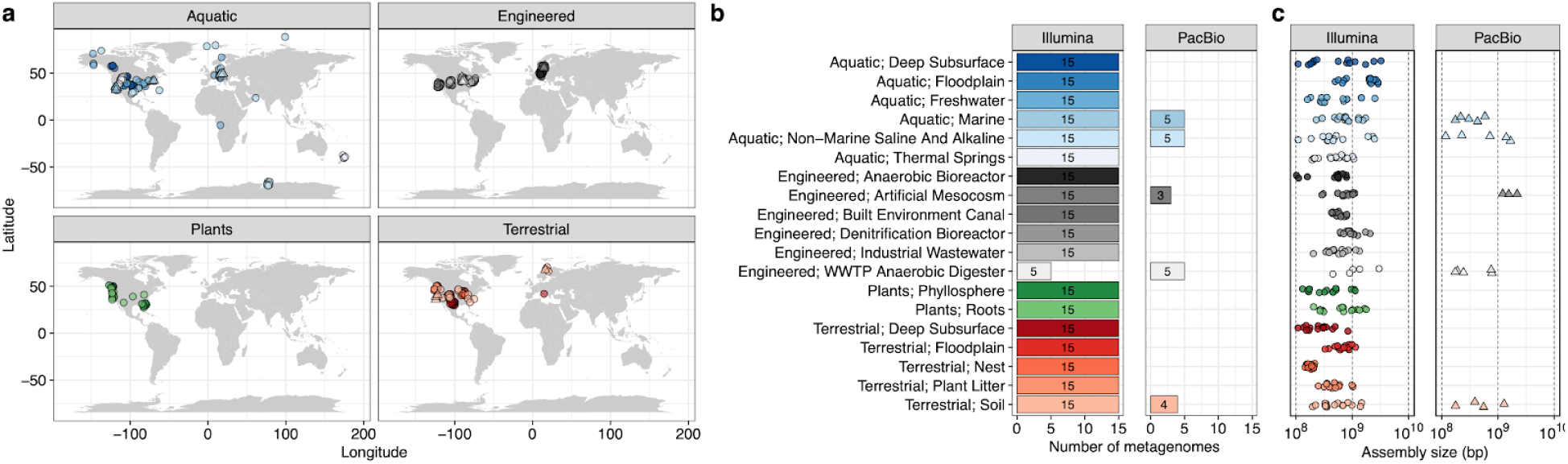
Global distribution and environmental diversity of JGI metagenomes selected for benchmarking. **(a)** Geographic distribution of the sampling locations for the metagenomes used to benchmark binning performance. Samples are faceted by four major ecosystem categories (biomes): Aquatic, Engineered, Plants, and Terrestrial. Map points indicate the latitude and longitude of sample collection. **(b)** Environmental subtype (sub-biome) and sequencing platform. The left column displays data for Illumina-sequenced samples, and the right column displays data for PacBio-sequenced samples. For each sub-biome (y-axis), the horizontal bar charts indicate the number of metagenomes included. **(c)** Dot plots represent the distribution of assembly sizes (in base pairs).

Furthermore, the selected metagenomes exhibit substantial variation in complexity, with assembly sizes ranging from approximately 100 Mb to over 3.5 Gb (Supplementary Figure 2c). This broad distribution of assembly sizes and environmental origins provides a rigorous, high-complexity testbed for evaluating the scalability and precision of binning algorithms under real-world conditions.

### 3.4. JGI bacterial isolates define a high-fidelity baseline for genomic reconstruction

To empirically determine rigorous quality thresholds for MAG recovery, we established a baseline of “gold standard” genomic fidelity using a reference set of bacterial isolates sequenced and curated by the Joint Genome Institute (JGI). We analyzed 15,554 JGI bacterial isolates that had undergone extensive Quality Assurance and Quality Control (QA/QC) to assess their completeness and contamination profiles using `CheckM2`. This analysis shows that high-quality reference isolates exhibit near-perfect completeness and negligible contamination. The mean quality metrics for this dataset were calculated at 99.82% completeness and 0.66% contamination (Supplementary Figure 3). These values situate the average isolate well within the ultra-high-quality spectrum (Contamination < 1%). This empirical baseline shows that while maximizing completeness is critical, minimizing contamination to sub-1% levels is equally necessary to approximate the fidelity of isolate-derived genomes for uses such as training robust genomic foundation models.

**Supplementary Figure 3.**
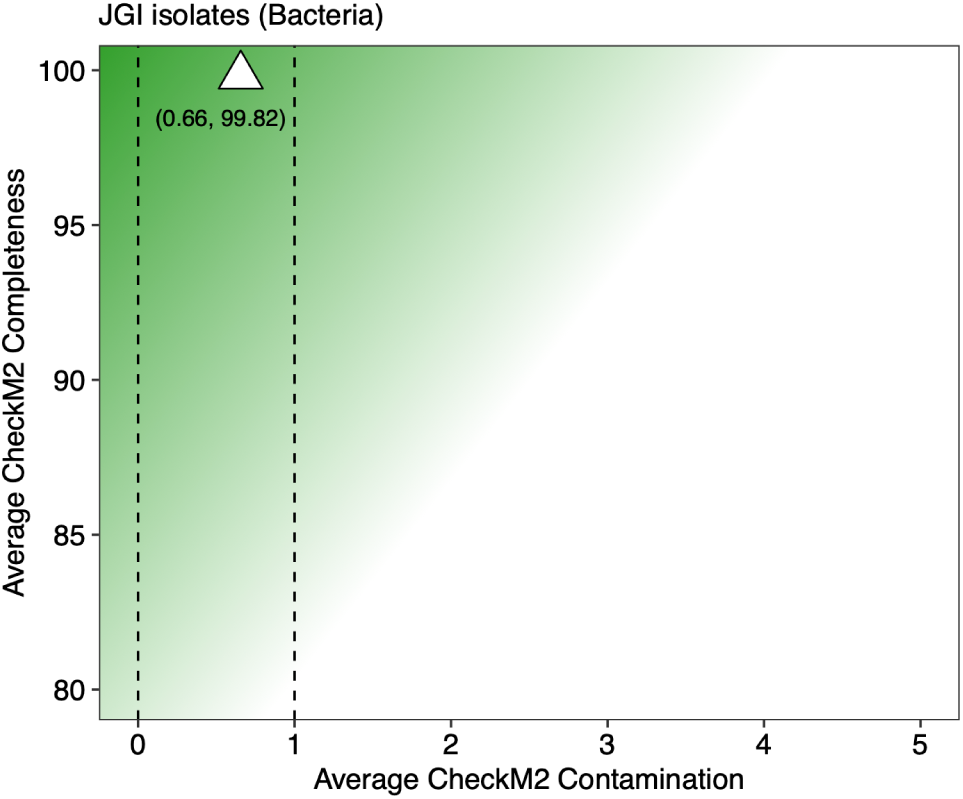
Quality baseline established by JGI bacterial isolates. Assessment of genomic quality metrics for a reference set of 15,554 JGI bacterial isolates. The plot displays the average completeness (y-axis) versus average contamination (x-axis) as assessed by `CheckM2`. The white triangle indicates the centroid of the distribution for the dataset, located at coordinates (0.66, 99.82). Vertical dashed lines mark the 0% and 1% contamination thresholds. The green gradient background serves as a visual guide for genomic quality, with darker green indicating higher completeness and lower contamination.

### 3.5. QuickBin delivers superior scalability and maximizes high-fidelity genome recovery

We evaluated the performance of `QuickBin` against four leading binners: `VAMB`, `SemiBin2`, `MetaBAT2`, and `COMEBin`; across the 297 diverse metagenomes curated for this benchmark (Supplementary Figure 2). `QuickBin` was run with default parameters and in ultra-loose (UL) mode. `SemiBin2` was run with default parameters using the global environment option and in self-supervised (SS) mode. All other binners were run with default parameters. Details on samples used in this benchmarking can be found in Supplementary Table S1. The assessment focused on two critical dimensions. First, the ability to successfully process large-scale data (scalability), and second, the total yield of high-quality genomes. Binners that allowed for GPU acceleration (`COMEBin`, `SemiBin2`, and `VAMB`) were provided with NVIDIA A100 40GB GPU nodes. The maximum allocation time was set to 24 hours (equivalent to more than 2,000× the median `QuickBin` runtime on these samples); and maximum RAM was set to 128GB (16× the `QuickBin` memory allocation).

We observed that the `MetaBAT2` (traditional CPU-based binning tool), `VAMB` (GPU-accelerated), and `QuickBin` (CPU-based), demonstrated higher robustness than deep-learning approaches under this computational resource availability (Figure 2a). While `MetaBAT2`, `VAMB`, and `QuickBin` successfully completed binning for all 297 samples (100% success rate), the other binners exhibited important failure rates due to runtime timeouts or memory exhaustion (Figure 2b). Specifically, `SemiBin2` failed to process 31 samples (10% failure rate), while `COMEBin` failed on 129 samples (43% failure rate), rendering it unsuitable for large-scale, automated pipelines without extensive resource investment.

**Figure 2.**
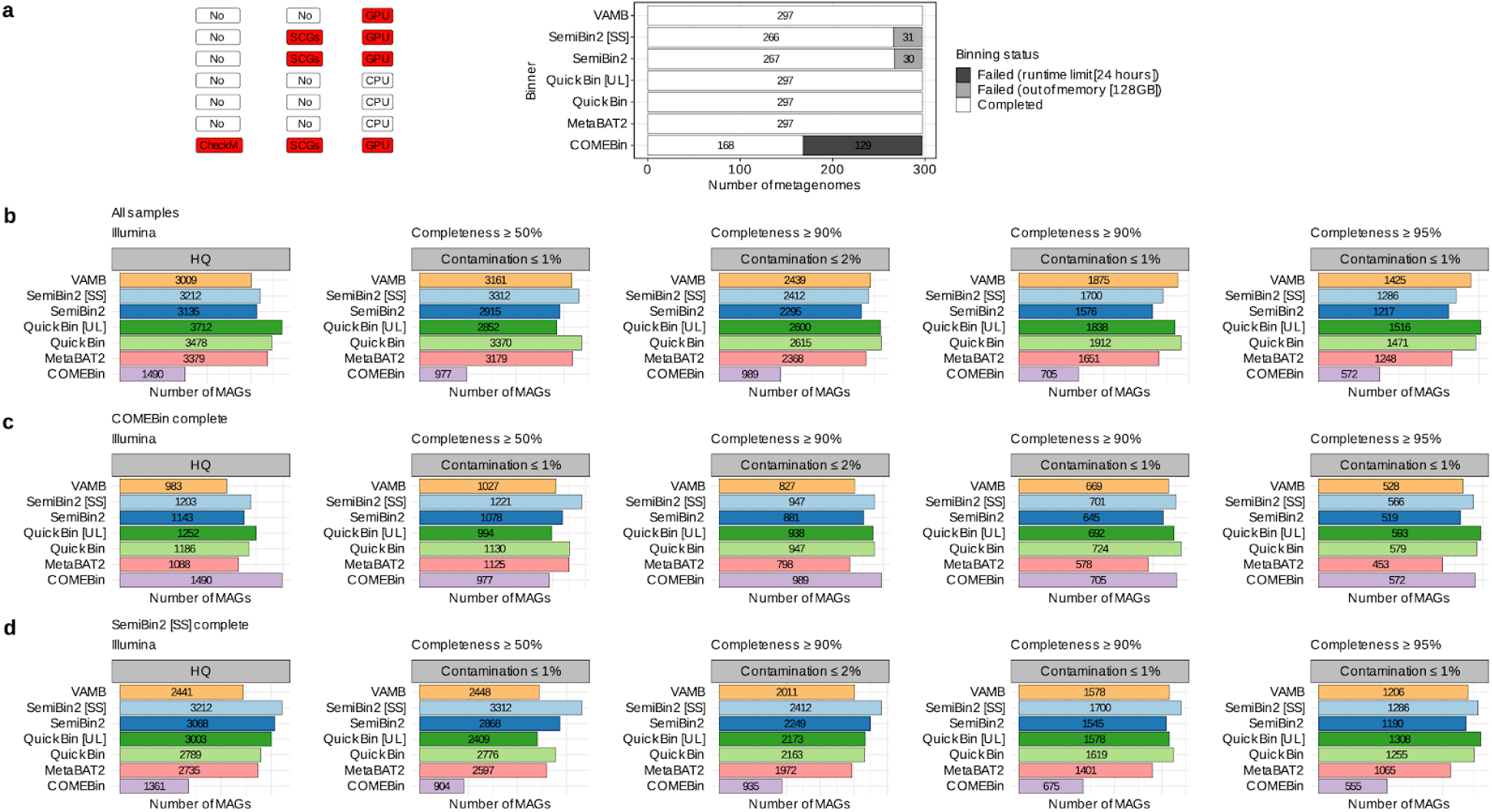
Scalability and genomic recovery performance of QuickBin versus state-of-the-art binners*. **(a)** Feature summary of the benchmarked binning tools, indicating dependency on `CheckM1`, dependency on Single-Copy Marker Genes (SCGs), and hardware requirements (CPU vs. GPU). Red boxes indicate resource dependency. The barplot shows the completion status for 297 diverse metagenomes. “Failed” indicates termination due to runtime limits or out-of-memory errors. `QuickBin`, `MetaBAT2`, and `VAMB` completed all samples, whereas `COMEBin` and `SemiBin2` experienced failure rates. **\*** Among the evaluated binners, `COMEBin` is the only method that does not provide a user-configurable minimum contig length for binning. Consequently, `COMEBin` was run with its default settings, whereas all other binners were run using a 1.5 kbp minimum contig length. **(b)** Total number of MAGs recovered from the full dataset (*n* = 297) across varying quality thresholds. `QuickBin` [UL]: ultraloose mode; `SemiBin2` [SS]: self-supervised mode. **(c)** Performance comparison on the specific subsets of metagenomes where `COMEBin` successfully completed processing. **(d)** Performance comparison on the specific subsets of metagenomes where `SemiBin2` successfully completed processing.

`QuickBin`‘s robustness advantage directly translated into superior overall genomic yield. When assessing the recovery of MAGs across all processed samples, `QuickBin` (specifically the `QuickBin` [UL] variant) consistently outperformed all competitors. Most notably, in the category of high completeness (≥95%) and ultra-low contamination (≤1%), `QuickBin` [UL] (ultraloose) recovered 1,516 MAGs, surpassing `SemiBin2` (1,286 MAGs), `VAMB` (1,425 MAGs), and `MetaBAT2` (1,248 MAGs) (Figure 2b). Even when restricting the analysis to the subsets of easy-to-bin samples where the resource-heavy tools successfully ran (“`COMEBin` complete” and “`SemiBin2` complete” subsets), `QuickBin` remained highly competitive, recovering higher numbers of high-fidelity MAGs; without requiring GPU acceleration or Single-Copy Marker Genes (SCGs) as in the case of `COMEBin` (Figure 2c) and `SemiBin2` (Figure 2d); or relying on `CheckM1` to pre-select the binning outcome as is the case for `COMEBin` (Figure 2c). While the `QuickBin` [UL] variant demonstrated the highest stotal MAG recovery in these benchmarks, the `QuickBin` default settings are engineered to prioritize absolute bin purity. This distinction is particularly relevant for downstream applications where even trace contamination (at or below the 1% sensitivity threshold of current marker-based tools) could propagate systematic errors.

### 3.6. Biome-specific performance

We next examined how the completeness distribution of recovered MAGs varies across biomes and binners (Supplementary Figure 4) under a contamination cutoff ≤1%. Across samples from Aquatic, Engineered, Plants and Terrestrial biomes, `QuickBin` produced MAG sets with higher completeness profiles, including MAGs in the highest completeness tier (≥95%) (Supplementary Figure 4a). In the `COMEBin`-complete subset (Supplementary Figure 4b), highly complete MAG counts in all biomes showed that `QuickBin` performed equally or better than other binners overall, with `SemiBin2` [SS] slightly leading in Engineered biomes. Similarly, the `SemiBin`-complete subset (Supplementary Figure 4c) mostly mirrors the all-samples patterns; but shows that `SemiBin2` [SS] performed better at Plant biomes. Overall, `QuickBin` performed equally or better than other binners.

**Supplementary Figure 4.**
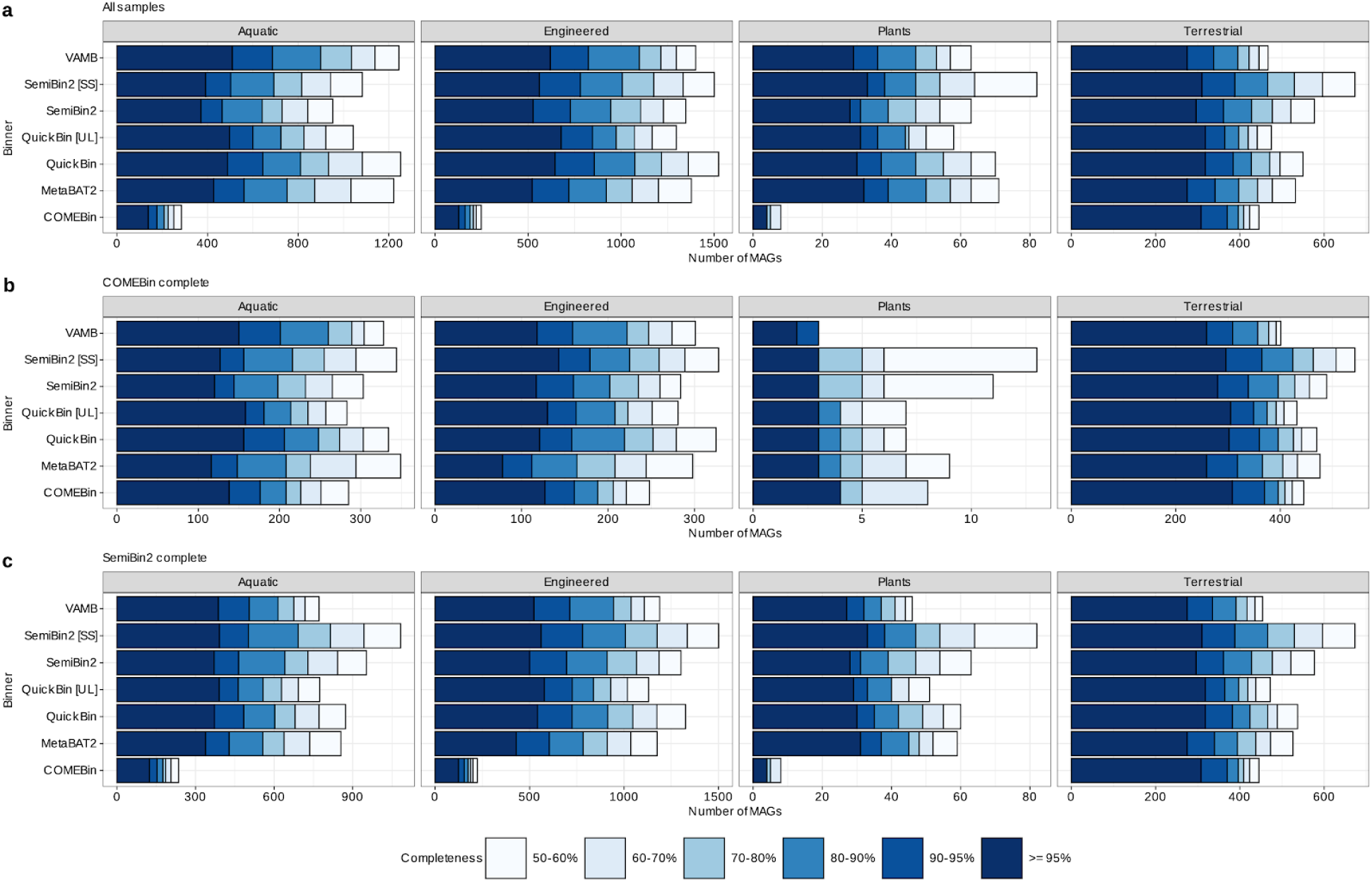
Biome-stratified completeness distributions across binners and completion-matched subsets. **(a)** Stacked distributions of MAG counts across completeness bins (50-60%, 60-70%, 70-80%, 80-90%, 90-95%, and ≥95%) and contamination ≤1% for each binner within Aquatic, Engineered, Plants, and Terrestrial biomes. **(b)** The same stratification restricted to the subset of metagenomes where `COMEBin` completed (“`COMEBin` complete”). **(c)** The same stratification restricted to the subset of metagenomes where `SemiBin2` completed (“`SemiBin` complete”).

### 3.7. Benchmarking QuickBin on synthetic data using CheckM2 as evaluator

Given that `CheckM2` is a learned estimator and may compress differences (Chklovski et al. 2023), and to further quantify performance under controlled ground truth, we benchmarked `QuickBin` against other binners on two synthetic datasets: a synthetic small community (*n* = 30 genomes) and a synthetic big community (*n* = 1,000 genomes). For each, we evaluated both a single-coverage design (one coverage file) and a multi-coverage design (multiple coverage files), and summarized recovery as the number of MAGs passing increasingly stringent QC thresholds according to `CheckM2` (Chklovski et al. 2023), culminating in our primary high-fidelity target (completeness ≥95% and contamination ≤1%) (Figure 3a-d). In the synthetic small, single-coverage benchmark (Figure 3a), `QuickBin` and `SemiBin2` recovered 19/30 MAGs at ≥95% completeness and ≤1% contamination, exceeding `VAMB`, `MetaBAT2` and `COMEBin` (Figure 3a). In the synthetic small, multi-coverage benchmark (Figure 3b), `QuickBin` and `QuickBin` [UL] achieved the highest recovery at the strictest tier (22/30 at ≥95% completeness and ≤1% contamination), outperforming all the other binners. Performance differences became more pronounced at scale. In the synthetic big, single-coverage benchmark (Figure 3c), `QuickBin` and `COMEBin` produced more MAGs across all QC tiers. In the synthetic big, multi-coverage benchmark (Figure 3d), `QuickBin` again performed the best. Across these synthetic data-based experiments, `QuickBin` consistently ranked at the top, for recovering near-complete, ultra-low-contamination genomes, with the `QuickBin` [UL] configuration providing an additional gain specifically in the ≥95% / ≤1% tier in both large benchmarks (Figure 3c,d).

**Figure 3.**
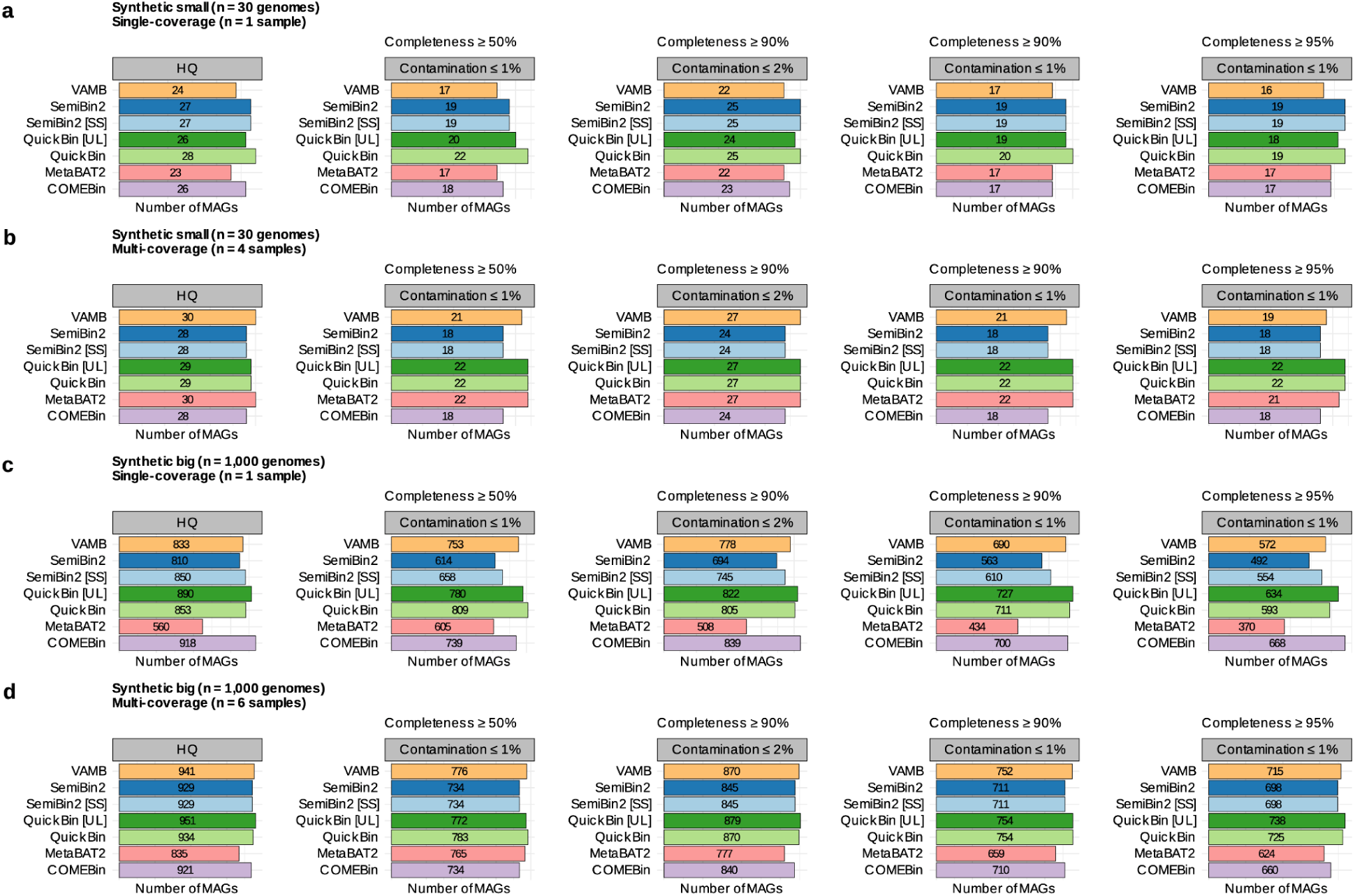
Synthetic benchmarking of binners under increasing QC stringency. **(a)** Synthetic small community (*n* = 30 genomes) under single-coverage (*n* = 1 sample). **(b)** Synthetic small community (*n* = 30 genomes) under multi-coverage (*n* = 4 samples). **(c)** Synthetic big community (*n* = 1,000 genomes) under single-coverage (*n* = 1 sample). **(d)** Synthetic big community (*n* = 1,000 genomes) under multi-coverage (*n* = 6 samples). Each panel reports the number of recovered MAGs for each binner under: HQ (>90% completeness and <5% contamination), completeness ≥50% with contamination ≤1%, completeness ≥90% with contamination ≤2%, completeness ≥90% with contamination ≤1%, and completeness ≥95% with contamination ≤1%.

### 3.8. Benchmarking QuickBin on synthetic data using ground truth as evaluator

To complement marker-based evaluations (Figure 3), we evaluated binner outputs against synthetic ground truth (see Methods for details) by tracking the origin of contigs assigned to each final bin and summarizing performance with four mapping-derived metrics: Genomes Represented (%), Sequence Recovery (%), Clean Bin Bases (%), and a composite Total Score (Figure 4a-d, see Methods for definition of Total Score). In this analysis, genomes represented captures how broadly a binner recovers the truth genomes present, sequence recovery summarizes how much true sequence is recovered into bins, and Clean Bin Bases describes the fraction of assembled bases assigned to uncontaminated bins derived from a single genome. This metric effectively filters out noise and chimeras, revealing the volume of high-fidelity genomic data that can actually be trusted for downstream analysis. (Figure 4a-d).

**Figure 4.**
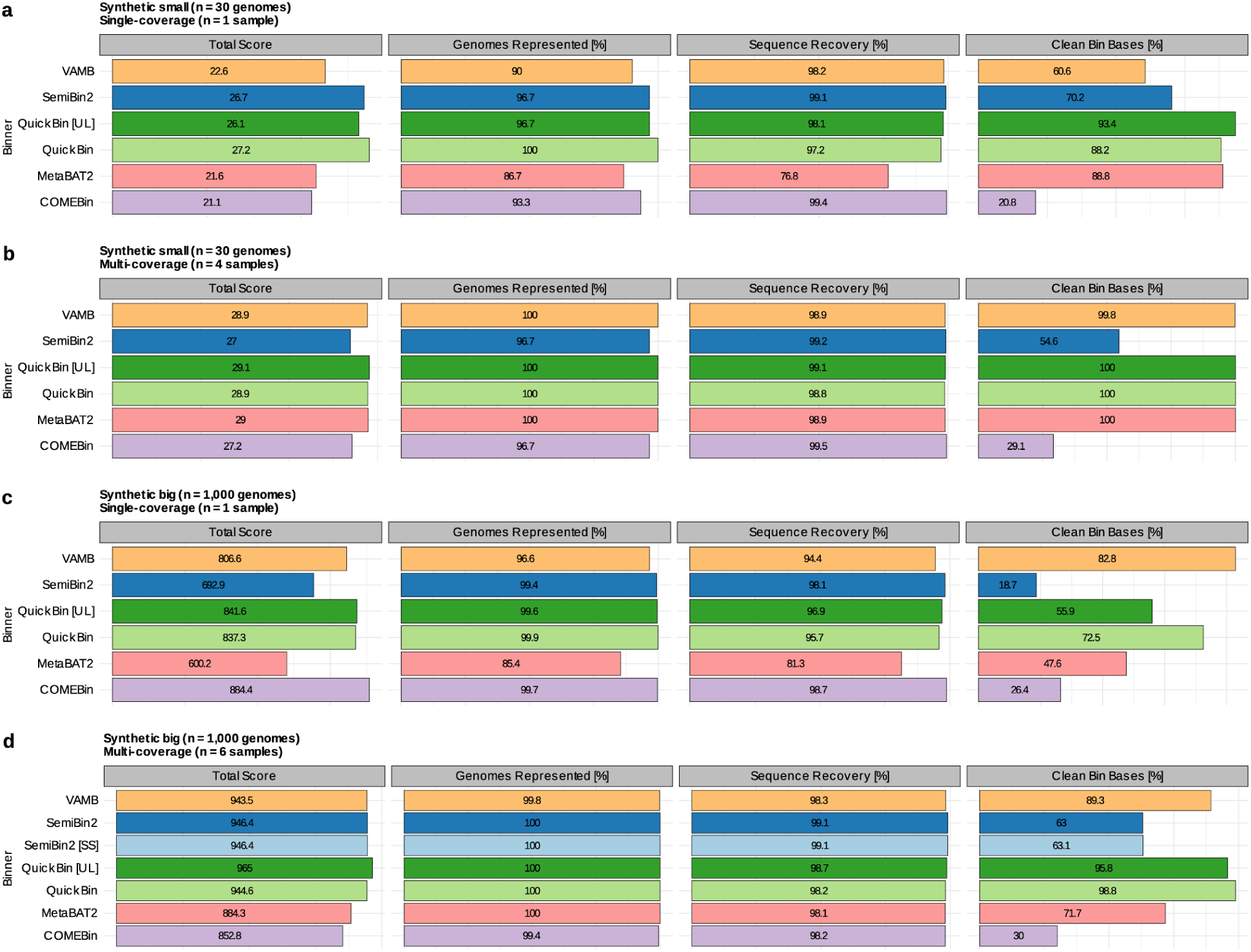
Ground truth evaluation of binners on synthetic communities by contig-origin mapping. **(a)** Synthetic small community (*n* = 30 genomes), single-coverage (*n* = 1 sample). **(b)** Synthetic small community (*n* = 30 genomes), multi-coverage (*n* = 4 samples). **(c)** Synthetic big community (*n* = 1,000 genomes), single-coverage (*n* = 1 sample). **(d)** Synthetic big community (*n* = 1,000 genomes), multi-coverage (*n* = 6 samples). For each binner, performance is summarized using mapping-derived metrics: Total Score [∑max(0,(completeness - 5*(contamination))^2^ , summed over all bins], Genomes Represented (%), Sequence Recovery (%), and Clean Bin Bases (%).

In the synthetic small, single-coverage benchmark (Figure 4a), `QuickBin` achieved the highest Total Score and genomes represented (100%) while maintaining high Sequence Recovery (97.2%). At the same time, `QuickBin` produced a high fraction of Clean Bin Bases (88.2%), closely matching `MetaBAT2` (88.8%) and exceeded by `QuickBin` [UL] (93.4%) (Figure 4a). In contrast, `COMEBin` recovered nearly all sequences (99.4%) and represented most genomes (93.3%), but had substantially lower Clean Bin Bases (20.8%), indicating extensive mixing of sources within binned sequence under ground truth (Figure 4a).

In the synthetic small, multi-coverage benchmark (Figure 4b), `QuickBin, QuickBin` and `QuickBin` [UL] recovered the full community with strong purity: both reached 100% genomes represented and 100% Clean Bin Bases, with near-complete sequence recovery (98.8 - 99.1%) (Figure 4b). `VAMB` and `MetaBAT2` also represented all genomes with high purity signal, while `SemiBin2` and `COMEBin` showed a larger drop in Clean Bin Bases (54,6% and 29.1%; respectively) despite high sequence recovery (Figure 4b). The Total Score ordering in this setting reflects these purity differences, with `QuickBin` [UL] highest (Figure 4b).

Performance differences became stronger in the synthetic big community. In the single-coverage case (Figure 4c), `COMEBin` achieved the highest total score at the expense of the lowest Clean Bin Bases (26.4%). `QuickBin` and `QuickBin` [UL] achieved the second highest Total Scores in this setting, supported by the highest Genomes Represented percentages, with the second highest Clean Bins Bases (72.4%) obtained by `QuickBin`; while the best purity was achieved by `VAMB` (82.8% Clean Bin Bases; Figure 4c). Between the two `QuickBin` modes, `QuickBin` [UL] achieved the highest total score in this setting (846.1), while `QuickBin` default retained the highest clean-bin fraction (Figure 4c), showing that the intended two `QuickBin` configurations prioritize slightly different balances between recovery and purity under ground truth. In the multi-coverage case (Figure 4d), `QuickBin` achieved complete recovery with exceptionally high purity by achieving 100% genomes represented, high sequence recovery, and the highest Clean Bin Bases among all methods (Figure 4d).

It is important to note that the divergence between marker-based estimates (Figure 3) and ground-truth mapping (Figure 4) reveals a critical blind spot in current metagenomic validation. While marker-based tools like `CheckM2` provide an accessible proxy for quality, their inherent resolution limits, specifically the inability to distinguish between a small percentage of contamination and 0% contamination, can mask significant chimeric artifacts. Our synthetic benchmarks demonstrate that methods optimized to maximize marker-based scores (e.g., `COMEBin` and `SemiBin2`) often achieve high completeness at the cost of destructive purity loss, with ground-truth Clean Bin Bases dropping as low as 18.7%. By contrast, marker-free methods, including `QuickBin`, `VAMB` and `MetaBAT2`, outperform marker-based methods in that metric whenever they can be directly compared. These results argue that synthetic datasets with known ground truth are the only reliable metric for validating high-fidelity binning algorithms, as relying on real-world datasets with unknown truth risks institutionalizing flawed outcomes as a baseline for methods evaluation.

Altogether, the ground-truth mapping analysis shows that `QuickBin` consistently recovers a large fraction of genomes and sequence while maintaining a high fraction of Clean Bin Bases, with the clearest advantage in the large multi-coverage setting where `QuickBin` achieves near-complete recovery together with near-complete bin purity (Figure 4d). These properties directly support `QuickBin`‘s goal of high-fidelity MAG reconstruction and help explain its strong recovery of near-complete, low-contamination genomes in the synthetic `CheckM2` benchmarks (Figure 3 and Figure 4).

### 3.9. Benchmarking on PacBio metagenomes

Although `QuickBin` was developed and optimized around short-read metagenome workflows, we benchmarked it on a PacBio metagenome dataset to provide users with a fair, decision-relevant comparison against other binners under long-read conditions (Supplementary Figure 5 and Supplementary Figure 6). We summarize results using the same increasingly stringent QC tiers used elsewhere in the study, culminating in the high-fidelity target of completeness ≥95% and contamination ≤1% (Supplementary Figure 5a). Across PacBio assemblies, `QuickBin` recovered fewer MAGs than other binners at every QC tier, consistent with the fact that `QuickBin` was not tuned for this data type (Supplementary Figure 5a). At the strictest tier, `QuickBin` produced 76 MAGs at completeness ≥95% and contamination ≤1% (`QuickBin` [UL], 62), compared to 206 for `SemiBin2`, 160 for `MetaBAT2`, 147 for `COMEBin`, and 142 for `VAMB` (Supplementary Figure 5a). Importantly, `QuickBin` still yielded a non-trivial set of near-complete, ultra-low-contamination MAGs on PacBio, indicating that its high-fidelity objective remains attainable even outside its primary optimization regime, albeit at reduced yield relative to the strongest PacBio performers. In this PacBio benchmark, the counts are unchanged between the all-samples and completion-matched panels because all binners completed for these datasets (Supplementary Figure 5a-c).

**Supplementary Figure 5.**
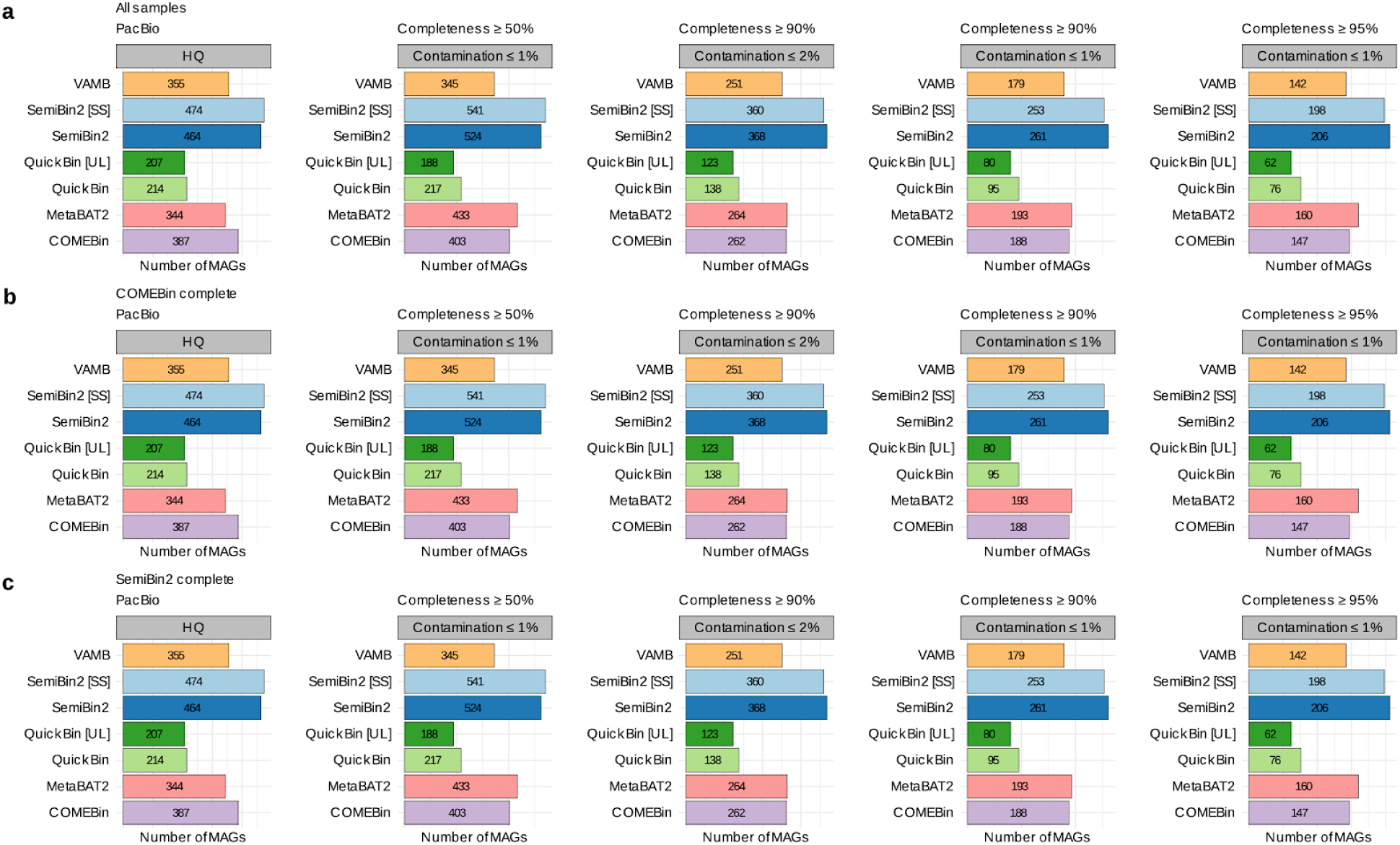
PacBio benchmarking across QC tiers. **(a)** PacBio metagenomes, all samples. Bar plots report the number of recovered MAGs per binner passing increasingly stringent QC tiers: HQ (completeness >90% with contamination <5%), completeness ≥50% with contamination ≤1%, completeness ≥90% with contamination ≤2%, completeness ≥90% with contamination ≤1%, and completeness ≥95% with contamination ≤1%. **(b)** The same analysis is restricted to the subset of metagenomes where `COMEBin` completed (“`COMEBin` complete”). **(c)** The same analysis restricted to the subset of metagenomes where `SemiBin2` completed (“`SemiBin2` complete”).

**Supplementary Figure 6.**
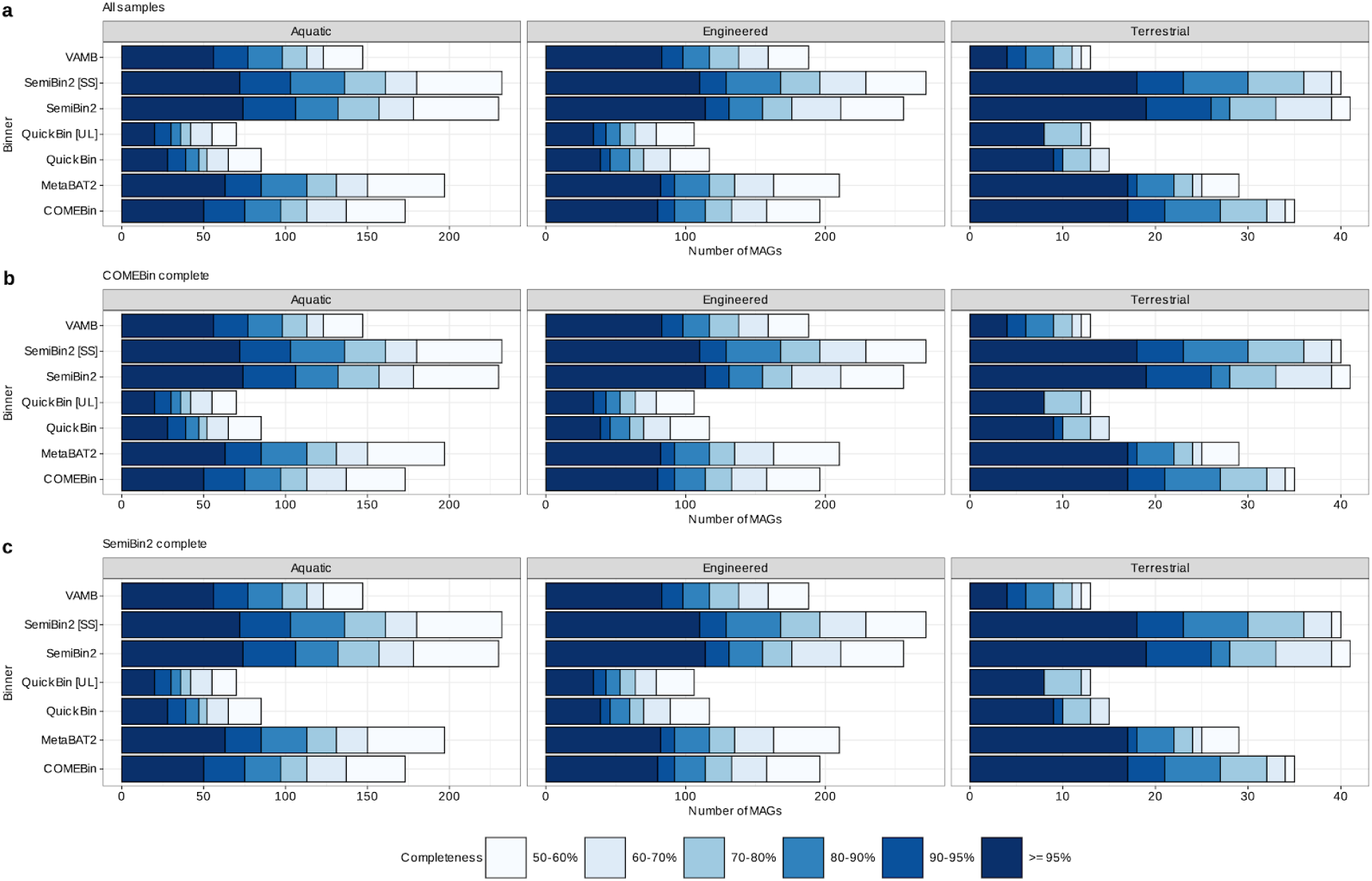
Biome-stratified completeness distributions for PacBio MAGs. **(a)** Distributions of MAG counts across completeness bins (50-60%, 60-70%, 70-80%, 80-90%, 90-95%, and ≥95%) for each binner within Aquatic, Engineered, and Terrestrial PacBio metagenomes (x-axis, number of MAGs). **(b)** The same stratification restricted to metagenomes where `COMEBin` completed (“`COMEBin` complete”). **(c)** The same stratification restricted to metagenomes where `SemiBin2` completed (“`SemiBin` complete”). Among the evaluated binners, `COMEBin` is the only method that does not provide a user-configurable minimum contig length for binning. Consequently, `COMEBin` was run with its default settings, whereas all other binners were run using a 1.5 kbp minimum contig length.

We further examined biome-stratified completeness distributions for PacBio assemblies across Aquatic, Engineered, and Terrestrial metagenomes (Supplementary Figure 6a-c). `QuickBin` contributed MAGs across completeness bins in each biome, but with lower totals than other binners (Supplementary Figure 6a). The same biome-level patterns are preserved in the completion-matched subsets (Supplementary Figure 6b,c) as all binners completed for all these datasets. In summary, these PacBio benchmarks provide a transparent boundary condition for `QuickBin`, relative to leading alternatives, `QuickBin` is not the highest-yield tool on PacBio assemblies, but it still recovers a measurable set of high-fidelity MAGs at completeness ≥95% and contamination ≤1%. Including these results supports responsible method selection by clarifying where `QuickBin` performs best, and where users may prefer PacBio-optimized approaches depending on study goals and sequencing strategy.

### 3.10. QuickBin strengthens eukMAG recovery

To evaluate eukaryotic metagenome-assembled genome (eukMAGs) recovery, we summarized each binner’s eukMAG output using `EukCC` completeness and compared the number of eukMAGs recovered across completeness bins (Figure 5; Supplementary Figure 7). This analysis focuses on completeness as estimated by `EukCC` (Saary et al. 2020); contamination cutoff was set to <10% as there are no established standards yet for eukMAG quality control.

**Figure 5.**
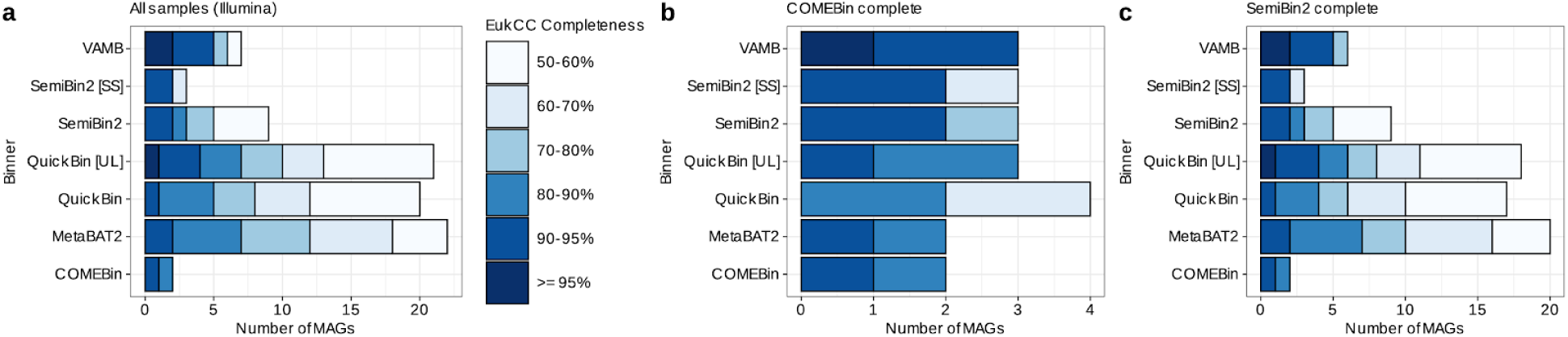
Eukaryotic MAG recovery on Illumina metagenomes assessed by EukCC completeness. **(a)** All Illumina samples. **(b)** Subset restricted to metagenomes where `COMEBin` completed (“`COMEBin` complete”). **(c)** Subset restricted to metagenomes where `SemiBin2` completed (“`SemiBin2` complete”). Horizontal stacked bars show the number of eukMAGs recovered by each binner at `EukCC`-estimated contamination <10%, stratified by `EukCC` completeness bins (color-coded): 50-60%, 60-70%, 70-80%, 80-90%, and 90-95%.

**Supplementary Figure 7.**
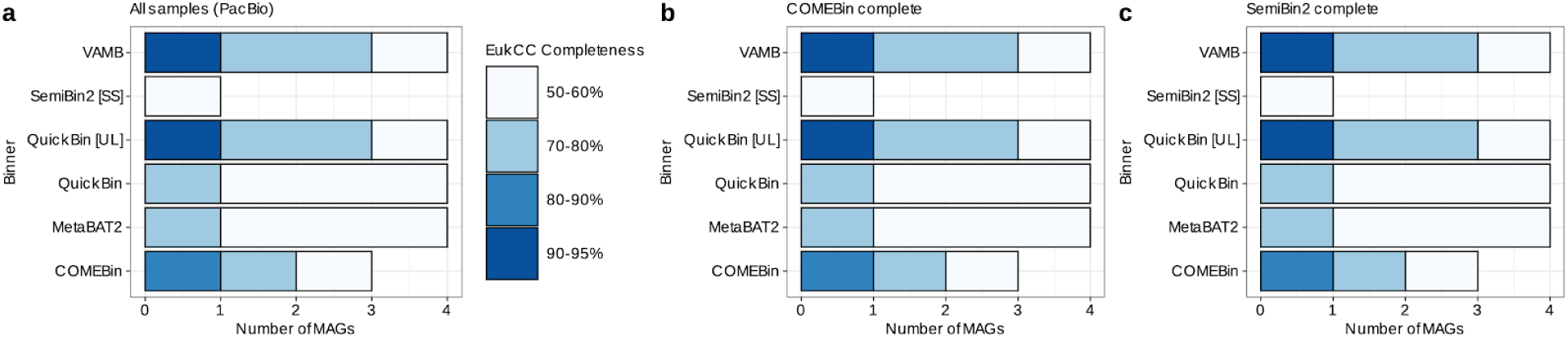
Eukaryotic MAG recovery on PacBio metagenomes assessed by EukCC completeness. **(a)** All PacBio samples. **(b)** Subset restricted to metagenomes where `COMEBin` completed (“`COMEBin` complete”). **(c)** Subset restricted to metagenomes where `SemiBin2` completed (“`SemiBin2` complete”). Horizontal stacked bars show the number of eukMAGs recovered by each binner at `EukCC`-estimated contamination <10%, stratified by `EukCC` completeness bins (color-coded): 50-60%, 70-80%, 80-90%, 90-95%, and ≥95%.

Across all Illumina samples, `QuickBin` and `MetaBAT2` recovered the largest eukMAG set among the tested binners and contributed eukMAGs across the full `EukCC` completeness spectrum, with particularly strong representation in the highest completeness bins shown at 80-90% and 90-95%; with only `QuickBin` and `VAMB` showing eukMAGs even at ≥95% (Figure 5a). In contrast, `COMEBin` and `SemiBin2` produced only a small number of eukMAGs in this benchmark, and their output was concentrated into fewer completeness strata (Figure 5a). To ensure that comparisons were not driven by differential completion, we repeated the same completeness stratification on completion-matched subsets. In the `COMEBin`-complete subset, the analysis is restricted to the metagenomes where `COMEBin` finished, which reduces the total number of eukMAGs available for all methods (Figure 5b). Within this matched subset, `QuickBin` remains among the top performers and continues to contribute eukMAGs in the higher completeness bins, with VAMB generating highly complete eukMAGs. In the complementary `SemiBin2`-complete subset, which restricted to metagenomes where `SemiBin2` completed, the overall structure of the result closely resembles the all-samples comparison, and `QuickBin`, `QuickBin` [UL], and `MetaBAT2` again recover the largest eukMAG totals, with substantial contributions in the highest completeness bins shown (Figure 5c). Together, panels a-c show that `QuickBin`’s strong eukMAG recovery on Illumina persists under completion-matched comparisons and is driven by increased recovery of higher-completeness eukMAGs rather than an isolated gain in low-completeness bins (Figure 5a-c).

We then performed the same `EukCC`-based stratification on PacBio metagenomes (Supplementary Figure 7a-c). Overall eukMAG yields are smaller in this PacBio evaluation, but `QuickBin` and `QuickBin` [UL] still recover eukMAGs across multiple completeness bins, including eukMAGs in the `EukCC` 90-95% bin (Supplementary Figure 7a). Completion-matched subsets (`COMEBin` complete and `SemiBin2` complete) preserve the same pattern as all PacBio samples completed successfully (Supplementary Figure 7b,c).

Overall, the `EukCC`-based benchmarking indicates that `QuickBin` is a strong option for eukaryotic MAG recovery, particularly in Illumina datasets where it yields the largest number of higher-completeness eukMAGs, and it remains competitive on PacBio where it continues to recover eukMAGs that reach `EukCC` high completeness (Figure 5; Supplementary Figure 7). This reflects its heritage as a taxonomically-unbiased clade-independent binning tool designed to cleanly separate prokaryotic and eukaryotic fractions. As in the labelled synthetic data contamination tests, marker-based binning approaches demonstrate severe weakness in eukaryotic bin recovery.

### 3.11. Computational efficiency enables deployable, scalable high-fidelity binning with QuickBin

Given that binning choices are often constrained as much by runtime and memory limits as by output quality, we quantified the computational footprint of each method under standardized benchmarking conditions. We therefore profiled wall-clock runtime and peak memory usage across Illumina and PacBio metagenomes, explicitly evaluating `QuickBin` in deployable CPU-only configurations (8 GB RAM with 8 CPU cores; 8 GB RAM with 32 CPU cores; and 32 GB RAM with 32 CPU cores) alongside `QuickBin` and other tools run under higher-resource availability settings (Figure 6; Supplementary Figure 8). It should be noted that QuickBin’s memory usage reflects the configurable nature of the Java Virtual Machine (JVM).

**Figure 6.**
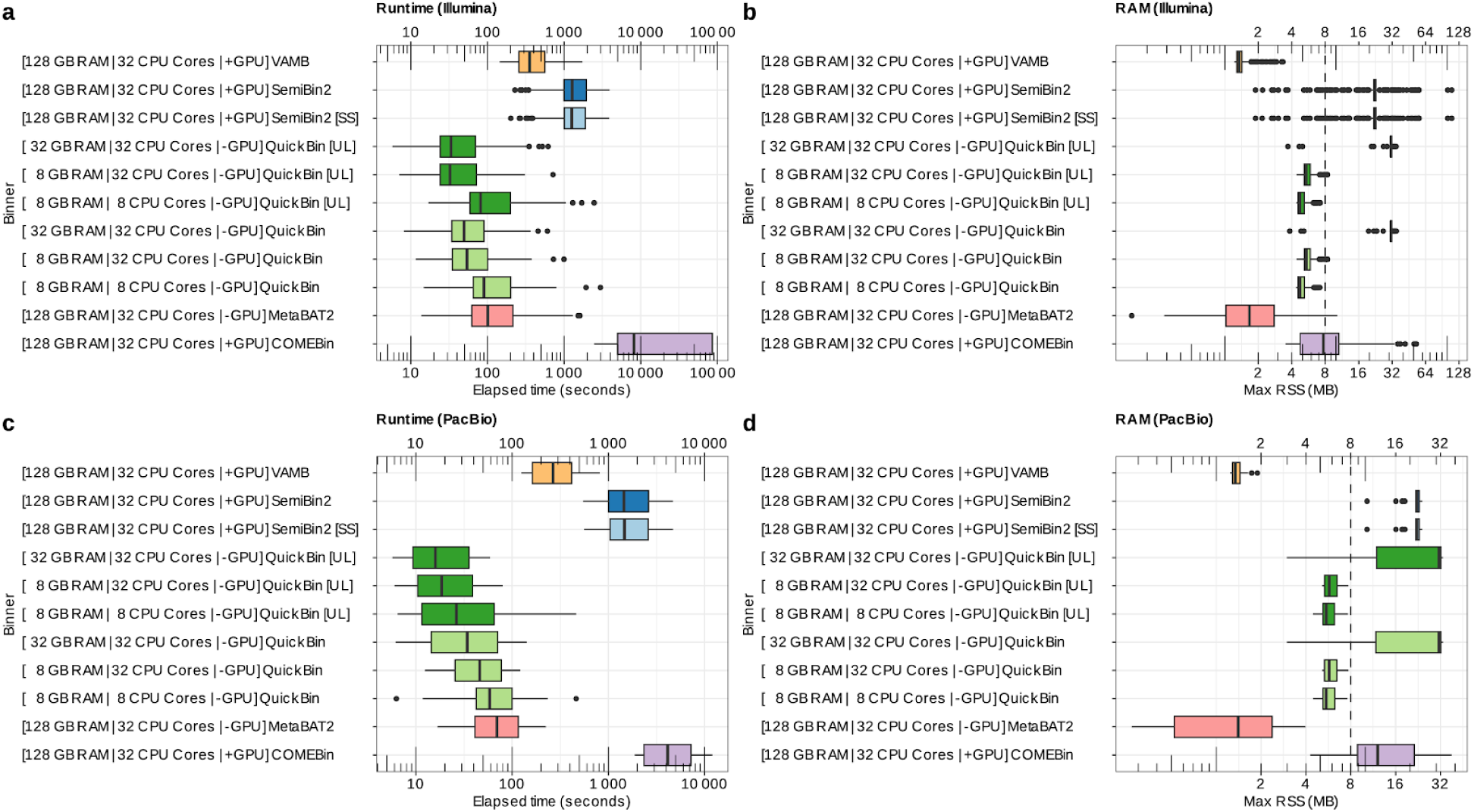
Runtime and peak memory usage across binners highlights QuickBin efficiency and deployability. **(a)** Runtime (Illumina). **(b)** RAM (Illumina). **(c)** Runtime (PacBio). **(d)** RAM (PacBio). Boxplots summarize per-metagenome elapsed time (seconds) and Max RSS (maximum resident set size, as labeled in the figure) for each binner and configuration. Labels indicate the hardware configuration used for each tool (RAM, CPU cores, and whether a GPU was used). `QuickBin` and `QuickBin` [UL] are shown under CPU-only configurations spanning 8 GB RAM (8 cores or 32 cores) and 32 GB RAM (32 cores), while several alternative methods are shown under 128 GB RAM and 32 cores, with GPU-acceleration indicated with “+GPU”, and CPU only runs with “-GPU”.

**Supplementary Figure 8.**
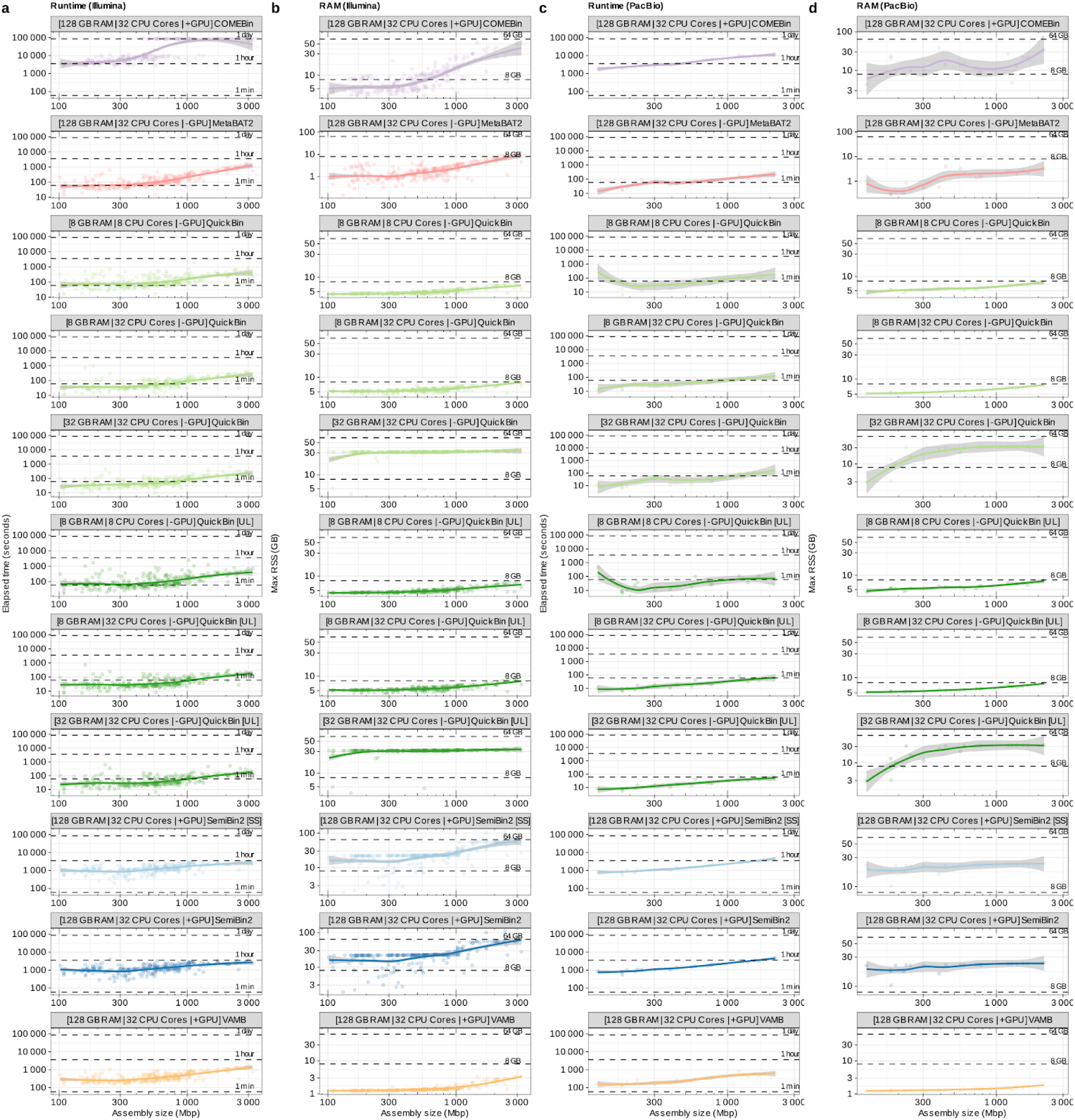
Scaling of runtime and memory with assembly size. **(a)** Runtime (Illumina) versus assembly size (Mbp). **(b)** RAM (Illumina) versus assembly size (Mbp). **(c)** Runtime (PacBio) versus assembly size (Mbp). **(d)** RAM (PacBio) versus assembly size (Mbp). Each row corresponds to one binner and configuration (as labeled). Points represent per-metagenome measurements; smoothed trend lines summarize scaling with assembly size. Runtime plots report elapsed time (seconds) and include dashed horizontal reference lines marking 1 min, 1 hour, and 1 day. Memory plots report Max RSS (GB) and include dashed horizontal reference lines at 8 GB and 64 GB.

Configurations provided with 8 GB of RAM use 8GB and those allowed 32 GB of RAM show higher peak RSS values, due to the JVM’s proactive allocation of available memory to optimize performance, not a strict requirement; Java will use the amount of memory you tell it to use.

On Illumina datasets, `QuickBin` showed a strong runtime advantage. Runtime distributions for all `QuickBin` configurations are within the tens of seconds range, whereas `COMEBin` shows longer runtimes in the range of days time (Figure 6a). Other GPU-accelerated binners (`SemiBin2`, `SemiBin2` [SS], and `VAMB`). Peak memory usage further supports deployability.

`QuickBin` maintains low Max RSS (Maximum Resident Set Size: the peak amount of a process’s physical RAM used) distributions across its configurations in the Illumina benchmark, consistent with practical execution under 8 GB RAM for most assemblies, while other methods include higher-memory requirements (Figure 6b). The same trends hold for PacBio datasets.

`QuickBin` remains the fastest method, dramatically separated from `COMEBin`, which again shows the longest runtimes (Figure 6c). Memory profiles on PacBio show that `QuickBin` can operate at low Max RSS in its 8 GB configurations, while higher-resource configurations (including `QuickBin` [UL] at 32 GB RAM) can increase peak memory for some samples, reflecting a tunable compute-throughput tradeoff (Figure 6d).

To examine scaling explicitly, we related runtime and memory to assembly size (Mbp). Across both Illumina and PacBio, `QuickBin` exhibits shallow scaling curves and remains within the most favorable time bands across the tested assembly-size range, whereas `COMEBin` increases steeply and trends toward substantially longer runtimes as assemblies grow (Supplementary Figure 8a,c). In memory scaling, `QuickBin` remains close to the 8 GB reference line in its constrained configurations, while several other methods trend upward with assembly size, approaching or exceeding higher memory bands (Supplementary Figure 8b,d).

To further assess scalability, we benchmarked `QuickBin` on an extremely large co-assembly from the “Terabase-Scale Coassembly of a Tropical Soil Microbiome” (Riley et al. 2023) project, generated from 3.4 Tbp of reads. The input comprised a 74-GB contigs file containing 55,342,847 contigs and 95 sorted read-alignment BAM files totaling 1.4 TB. `QuickBin` completed in 10 h 56 min 23 s using 64 threads with a peak memory usage of 383 GB, producing 5,028 bins.

Our computational efficiency benchmarks show that `QuickBin` delivers high-throughput binning with low peak memory and short runtimes across both Illumina and PacBio, and that it scales favorably with assembly size under CPU-only, low-RAM configurations (Figure 6; Supplementary Figure 8). When combined with `QuickBin`’s strong recovery of high-fidelity MAGs in the ≥95% completeness and ≤1% contamination regime shown in earlier benchmarks, these efficiency properties directly support deployable, large-scale reconstruction of near-complete, ultra-low-contamination genomes.

## 4. Discussion

Genome-resolved metagenomics continues to expand in scale and impact, but the field still faces a persistent methodological challenge pronounced by methods that maximize MAG yield do not always maximize MAG fidelity, and methods that achieve high accuracy can be difficult to deploy broadly due to compute and hardware requirements (Meyer et al. 2022; Han et al. 2025). This trade-off matters because many downstream uses, including comparative genomics, ecological inference, and the construction of curated genome resources, increasingly depend on consistently high-quality reconstructions rather than maximal genome counts (Bowers et al. 2017; Parks et al. 2022; Gurbich et al. 2023). In this study, we show that `QuickBin` addresses this gap by explicitly optimizing for the recovery of near-complete, ultra-low-contamination genomes while remaining deployable in resource-constrained settings, improving the practical accessibility of high-fidelity binning at scale.

In practice, `QuickBin` is intended to let metagenomics scientists run genome-resolved analyses at scale while still targeting stringent bin quality, especially bins meeting completeness at least 95% and contamination at most 1%. This matters because many projects require binning dozens to thousands of assemblies, often with single-coverage or multi-coverage designs, where compute limits and incomplete runs can force compromises in tool choice and QC stringency. By making conservative merging feasible under bounded resources (Figure 1), `QuickBin` is designed to reduce the burden of downstream manual triage and refinement by shifting effort toward a smaller set of higher-confidence bins that are more appropriate for comparative genomics and ecological inference.

Across real short-read metagenomes, QuickBin produced the highest yield of high-fidelity MAGs under the strictest target regime (completeness ≥95% and contamination ≤1%), with the QuickBin [UL] configuration recovering 1,516 genomes in this category (Figure 2b).

Importantly, this performance was always paired with operational robustness (Figure 2a) and superior computational performance (Figure 6a-b). `QuickBin`, `MetaBAT2`, and `VAMB` completed all Illumina assemblies, whereas SGC-dependent tools (`COMEBin` and `SemiBin2`) showed non-trivial failure rates on the same dataset, consistent with the “high compute, lower deployability” side of the dichotomy reported in recent large-scale benchmarks (Han et al. 2025). These results position `QuickBin` as a practical alternative to recent deep-learning and post-processing frameworks (`SemiBin2`, `COMEBin`, `MetaBinner`), particularly when the goal is to assemble a high-confidence genome set rather than to maximize bin count by optimizing for non-perfect grading systems like `CheckM2` (Pan et al. 2022, 2023; Wang et al. 2023, 2024). QuickBin, by design, does not bias binning or discovery as it does not use any SCG information, and it is CheckM agnostic.

`QuickBin` achieved top performance always in the regime of ≥95% completeness and ≤1% contamination in both real metagenomes (Figure 2) and synthetic communities (Figure 3). A key contribution of this work is that `QuickBin`’s advantage at high stringency is not limited to marker-based QC (Figure 4). Interestingly, our ground-truth-derived analyses show that high sequence recovery can coexist with substantial impurity for some methods under certain settings, reinforcing the value of combining marker-based QC (for example `CheckM2`) (Chklovski et al. 2023) with orthogonal ground-truth or proxy purity assessments whenever feasible (Meyer et al. 2022). The contrast of `QuickBin` with `COMEBin` in the ground-truth-derived analyses settings is instructive; `COMEBin` recovered nearly all sequences and most genomes, yet produced only 21-30% Clean Bin Bases, indicating extensive mixing despite high apparent recovery (Figure 4). Our findings demonstrate that `QuickBin` narrows the quality gap between classic coverage-and-composition binners (for example `MetaBAT2`) and more compute-intensive modern approaches (for example `SemiBin2`, `COMEBin`), especially in the regime most aligned, although stricter, with MIMAG high-quality expectations and high-impact downstream use (Bowers et al. 2017; Kang et al. 2019; Nissen et al. 2021; Pan et al. 2023; Wang et al. 2024; Chklovski et al. 2023). In parallel, `QuickBin`’s strong recovery of eukaryotic MAGs under `EukCC` (Figure 5) shows it can contribute to genome-resolved efforts beyond bacteria and archaea, an area of growing interest given the emerging role of eukaryotic MAG catalogs in ecology and evolution (Massana and López-Escardó 2022).

As expected from its design focus on short-read workflows, `QuickBin` did not lead in overall MAG yield on PacBio assemblies. Nevertheless, `QuickBin` recovered a set of high-fidelity PacBio MAGs, showing that the high-fidelity objective remains attainable outside its primary optimization objective (Supplementary Figure 5). This result is relevant in light of rapid progress in long-read metagenomic assembly and analysis (for example HiFi-driven improvements in genome resolution and new long-read assemblers), which is reshaping expectations for contiguity and strain resolution (Kim et al. 2022; Bouras et al. 2024; Sereika et al. 2025).

`LorBin` (Xue et al. 2025), a new long-read metagenome binning tool, was published after we completed our benchmarking and therefore was not included in our comparisons, but it highlights an exciting direction for long-read metagenome binning as these approaches mature for new long-read based compendiums (Sereika et al. 2025). Further work on `QuickBin` development will be focused on also optimizing for long-read technologies.

A central practical outcome of this study is that `QuickBin`’s high-fidelity performance is paired with clear computational advantages. Figure 6 shows that `QuickBin` runs in CPU-only configurations and remains functional under memory-constrained settings (including 8 GB RAM), while the strongest deep-learning methods in this comparison were evaluated under GPU-enabled, high-memory configurations. These characteristics directly address deployability bottlenecks emphasized in recent community assessments, where method performance must be considered alongside throughput, hardware accessibility, and failure modes when benchmarking at realistic scales (Meyer et al. 2022; Han et al. 2025). In practice, the combination of high-fidelity recovery and low-resource execution enables `QuickBin` to function as a default high-confidence binner for large studies, including those that aim to build curated genome collections compatible with GTDB-based taxonomy and modern genome resource pipelines (Parks et al. 2022; Chaumeil et al. 2022; Parks et al. 2025; Gurbich et al. 2023).

Our results also have implications beyond traditional comparative genomics. Large, high-quality MAG datasets are increasingly used as training data for foundation models in biology, including nucleotide and protein language models (for example Nucleotide Transformer (Dalla-Torre et al. 2025), Evo (Nguyen et al. 2024), and ESM-class models (Lin et al. 2023)), which are sensitive to systematic noise and mislabeled content in their training sets. In that setting, high-fidelity binning is not only a quality-control concern but also a data-governance step that can influence what biological signals models learn. By preferentially recovering ≥95% complete and ≤1% contaminated genomes at scale, `QuickBin` can help shift genome-resolved metagenomics toward datasets that are better aligned with both rigorous biological inference and modern data-centric modeling.

Existing binning methodologies are compared based on, and thus often optimized for, MIMAG standards and marker-gene-based quality estimation. We have shown that this can lead to poor outcomes for both eukaryotic-fraction binning and the ability to produce truly contamination-free MAGs, due to the sparse distribution of marker genes used by existing tools and thus inherently low sensitivity to particularly shorter contaminant contigs. MIMAG itself, with a highest-quality category allowing 5% contamination, inflates benchmarks scored in “#HQ Bins recovered” by classifying pure bins together with bins too contaminated to be considered for release as an isolate. Benchmarks using synthetic data with a known ground truth are the only way to audit these estimators and expose failures (such as the 20.8% clean-base purity observed in marker-optimized tools) that are otherwise concealed by the inherent resolution limits of current marker-gene heuristics. New standards are needed to go beyond MIMAG and accurately evaluate binning of mixed communities, allowing the MAGs to safely be used alongside isolates for confident scientific discovery. Specifically, unification of grading bins across clades, and the presence of higher-stringency categories such as ≥95% completeness and <1% contamination would encourage the development and robust benchmarking of universal binning methodologies and the use of isolate-quality MAGs.

In summary, `QuickBin` provides a deployable path to high-fidelity genome reconstruction at scale. Across real Illumina metagenomes and controlled synthetic benchmarks, `QuickBin` consistently ranked at or near the top for recovering ≥95% complete and ≤1% contaminated genomes, while maintaining robust completion rates and substantially lower hardware demands than other tools. `QuickBin` delivers in a few seconds results that can require days on enterprise GPU nodes for competing methods, and memory requirements scale linearly rather than quadratically by avoiding all-to-all contig comparison. These properties make `QuickBin` well suited for large-scale, high-confidence genome catalog construction and for workflows where MAG fidelity, reproducibility, and compute accessibility are primary optimization objectives, which is increasingly the case in modern genome-resolved metagenomics.

## 5. Methods

### 5.1. Software used for benchmarking

Details on version, installation method, and running methods for software and databases used in this project can be found in Supplementary Table S3. In brief, software used in this study and corresponding versions were as follows: Binning was performed with `QuickBin` (BBTools 39.60; deployed via `docker` image `bryce911/bbtools:39.60`), `MetaBAT2` v2.18, `SemiBin2` v2.2.0, `COMEBin` v1.0.4, and `VAMB` v5.0.4; all binners and their respective dependencies, including `CUDA` libraries required for GPU usage, were installed via `pixi` unless otherwise noted. Quality assessment and filtering used `CheckM2` v1.1.0 (installed via `pixi`) and `EukCC` v2.1.3 (executed from the container quay.io/microbiome-informatics/eukcc:2.1.3), and `gradebins` from `BBTools` (v39.54; deployed via the `docker` image at `bryce911/bbtools:39.54`). Taxonomic assignment used `GTDB-Tk` v2.4.1 (installed via `mamba` environment). Read mapping processing used `samtools` v1.22.1 (installed via `pixi`). Reference databases included the `GTDB-Tk` data package for GTDB release r226 and the `EukCC` database v1.2, both retrieved via `wget`.

### 5.2. Metagenome datasets, assemblies, alignments, and reference databases

Metagenomes used for benchmarking (*n* = 297) were obtained from the IMG/M (Chen et al. 2023) through the JGI Data Portal (<u>data.jgi.doe.gov</u>), including assembled contigs and the corresponding read-alignment files (SAM/BAM) used for coverage profiling. Sample identifiers, project identifiers, sequencing platform labels (Illumina or PacBio), and biome assignments used throughout the manuscript are provided in Supplementary Table S4, enabling retrieval of the exact input datasets used for all comparisons. Biomes were selected and grouped into the four categories: Aquatic, Engineered, Plants, and Terrestrial (Supplementary Table S4). For real metagenomes, assemblies spanned a broad range of total assembled sequence, from 101,945,291 bp to 3,233,917,651 bp. Illumina metagenomes selected were only those assembled with `metaSPAdes` (Nurk et al. 2017), and PacBio metagenomes selected were only those assembled with `metaFlye` (Kolmogorov et al. 2020) (assembly method detailed per sample in Supplementary Table S4). All binners were run on the same assemblies and associated coverage inputs derived from these contigs and alignments, and downstream quality-control evaluations were applied uniformly across methods.

We selected real metagenomes using IMG metadata to construct a recent, public, and ecologically annotated benchmarking set suitable for consistent comparisons. Starting from restricting to metagenome analysis projects added on or after 1 January 2019 and required that sequencing was exclusively done by the DOE Joint Genome Institute as we know the through QA/QC these projects undergo. All datasets were unrestricted and of public availability. To standardize inputs, we limited assemblies to those generated with `metaSPAdes` or `metaFlye` and restricted sequencing platforms to Illumina or PacBio. We retained only samples with geographic coordinates (latitude and longitude) to supplement metadata with biome- and geography-aware information. To avoid atypical assemblies that complicate scaling comparisons, we constrained the total assembled sequence to 100 Mbp to 10 Gbp. Finally, we required non-unclassified ecosystem annotations to enable robust stratification into the biome and sub-biome categories used throughout the benchmarking.

Reference databases were version-pinned. For prokaryotic genome quality estimation we used the `CheckM2` reference database v1.1.0. For eukaryotic genome quality estimation, we used the `EukCC` database v1.2. For prokaryotic taxonomy, we used the `GTDB-Tk` data package corresponding to GTDB release r226.

### 5.3. Synthetic ground-truth data generation

To ensure rigorous training and validation of the neural network, we developed a novel data generation methodology designed to mimic the structural and topological artifacts of real metagenomic assemblies (e.g., breakpoints, collapsed repeats, and uneven coverage) rather than utilizing idealized “shredded” references that oversimplify the features of metagenomes. The pipeline follows these steps:

#### 5.3.1. Topological re-assembly

Reference genomes were individually re-assembled using Tadpole with an extended k-mer length (`k=155)` and permissive extension parameters (`mincountseed=1`, `mincountextend=1`). This step forces the assembler to traverse the de Bruijn graph for single-copy regions while naturally collapsing repetitive elements, thereby producing contigs with realistic graph topology and coverage spikes, rather than uniform linear fragments.

#### 5.3.2. Synthetic read generation

Error-free synthetic paired-end reads were generated from the original reference genomes using RandomReads (Bushnell et al. 2025) at 10× depth. Taxonomic identifiers (TaxIDs) were encoded into the source filenames to maintain provenance.

#### 5.3.3. Strict Alignment & Annotation

Synthetic reads were mapped back to the re-assembled contigs using `BBMap` (Bushnell 2014) in “semiperfect” mode (`semiperfectmode`, `ambig=random`). This alignment mode enforces perfect identity for the mapped portion of the read but permits soft-clipping at contig termini, ensuring that valid alignments are retained at breakpoints and contig ends where assemblers typically truncate.

#### 5.3.4. Ground-truth assignment

Contigs were annotated using ContigRenamer, a custom utility (`renamebymapping.sh`: https://bbmap.org/tools/renamebymapping) that parses alignment headers to calculate per-contig coverage and assigns the taxonomic ID via a majority-vote consensus of mapped reads. This ensures that chimeric contigs or collapsed repeats are labeled based on the dominant contributing organism, providing a realistic “ambiguous” class for the neural network to evaluate. The inputs input to the network are detailed in Table 1.

### 5.4. Computing environment and benchmarking resource caps

All benchmarks were executed on the NERSC Perlmutter supercomputer (https://www.nersc.gov/). Perlmutter CPU nodes are dual-socket systems with 2x AMD EPYC 7763 (Milan) CPUs, 64 cores per socket (128 physical cores per node), and 512 GB DDR4 memory. Perlmutter GPU nodes are single-socket systems with 1x AMD EPYC 7763 CPU (64 physical cores) and 256 GB DDR4 host memory, paired with 4x NVIDIA A100 GPUs; most GPU nodes provide 40 GB A100 GPUs, and a subset of GPU nodes provide 80 GB A100 GPUs.

Although Perlmutter nodes provide substantially more CPU cores and memory than required for many metagenome binning jobs, we enforced explicit per-run resource caps to ensure fair, deployability-relevant comparisons. For each benchmark run, we limited execution to the CPU core count and RAM specified for that tool and configuration (as reported in the corresponding Results sections and figure panels) by requesting those resources via Slurm and by setting tool thread counts accordingly. All jobs were submitted with a maximum wall-clock time of 24 hours (2,000× the median `QuickBin` runtime), and runs not completed within this limit were recorded as incomplete for the completion-rate analyses.

### 5.5. Binner parameterization

All binners were executed through a single orchestration layer to enforce uniform inputs, logging, and resource allocation across tools. Each run consumed an assembly FASTA and one sorted BAM file (single coverage) or a directory of sorted BAM files aligned to the same contig set (multi coverage). Binner outputs were collected as per bin FASTA files and standardized into a consistent directory layout for downstream QC and comparisons.

#### 5.5.1. Global thresholds

To make outputs directly comparable across methods, we applied the same reporting thresholds throughout benchmarking. We used a minimum contig length of 1,500 bp and a minimum bin size of 100,000 bp. Among the evaluated binners, `COMEBin` is the only method that does not provide a user-configurable minimum contig length for binning. Consequently, `COMEBin` was run with its default settings, whereas all other binners were run using a 1.5 kbp minimum contig length.

#### 5.5.2. CPU, memory, and runtime control

Each binner was run under explicit CPU and RAM allocations corresponding to the benchmark configuration described in the figure panels, and thread counts were set deterministically from those allocations. All runs were subject to a maximum wall clock time of 24 hours (2,000× the median `QuickBin` runtime) at the scheduler level. For `QuickBin` (written in Java), Java heap size was capped by a run specific memory budget.

#### 5.5.3. Binner specific settings

##### 5.5.3.1. QuickBin

`QuickBin` was run in the default mode and the ultra loose mode (“`QuickBin` [UL]”), as reported in figures. All `QuickBin` runs used SIMD acceleration. To standardize coverage inputs, `QuickBin` runs required sorted BAMs and regenerated coverage profiles every time from the provided alignments.

##### 5.5.3.2. MetaBAT2

To guarantee fairness and comparability with all other binners, MetaBAT2 was executed using the RunMetaBAT command which incorporates both the coverage computation and the MetaBAT2 binning.

##### 5.5.3.3. VAMB

`VAMB` was run using a fixed random seed for reproducibility. GPU mode was enabled.

##### 5.5.3.4. SemiBin2 and SemiBin2 [SS]

`SemiBin2` was run in two configurations, standard and self supervised (“SS”). The execution mode was selected based on whether coverage was single sample or multi sample, and GPU mode was enabled.

##### 5.5.3.5. COMEBin

COMEBin was run with GPU mode enabled.

### 5.6. Computational resources measurement

Computational resource consumption was quantified for each bioinformatics tool executed within our benchmarking pipeline using `GNU time` v1.9 (https://www.gnu.org/software/time/) invoked with the verbose flag (-v). This utility was automatically prepended to all tool invocations via wrapper functions to ensure consistent resource tracking across all binning algorithms and quality control steps. For each tool execution, `GNU time` captured the following metrics: user CPU time (seconds spent executing in user mode), system CPU time (seconds spent in kernel mode), elapsed wall-clock time (total real time from process start to completion), percent CPU utilization (ratio of CPU time to wall-clock time), and maximum resident set size (peak physical memory usage in kilobytes).

### 5.7. Abbreviations, short forms, and definitions

**MAG**: Metagenome-assembled genome, a draft genome reconstructed by binning contigs from a metagenome assembly into genome bins.

**eukMAG**: Eukaryotic metagenome-assembled genome, a MAG inferred to be eukaryotic and evaluated with eukaryote-focused QC.

**HQ**: High-quality MAG. In this manuscript, use an explicit threshold definition at first use (for example the MIMAG HQ definition, or your study-specific HQ tier).

**SCG**: Single-copy gene. Marker genes expected to occur once per genome, used for estimating completeness and contamination and for some binning/refinement approaches.

**VAE**: Variational autoencoder. A deep generative neural network used to learn low-dimensional representations of contigs from features such as k-mer composition and coverage; VAMB is an example of a VAE-based binner.

**SS (as in `SemiBin2` [SS])**: Self-supervised. A learning setting where the model trains using automatically generated labels or constraints rather than curated ground-truth labels.

**UL (as in QuickBin [UL]):** ultraloose mode of QuickBin.

**GC content (GC%)**: Fraction of bases in a sequence that are G or C, often used as a genome composition signal.

**N50**: A contiguity metric defined as the contig length L such that 50% of the total assembled sequence length is contained in contigs of length ≥ L.

**Max contig length**: The length of the longest contig in a MAG or assembly.

**CDS**: Coding DNA sequence, a predicted protein-coding gene region. In your analyses, “total CDS” is the number of predicted CDS per MAG.

**ANI**: Average nucleotide identity.

**RSS**: Resident set size. The amount of physical memory a process occupies; Max RSS is the peak resident memory during execution.

**Wall-clock time / elapsed time**: Actual real-world time taken for a program run, as opposed to CPU time.

**CPU core / thread**: Execution units allocated to a job; specify whether tools were run with multithreading and how threads were set.

**Coverage**: The average number of reads supporting each base or the mean read depth on a contig; in binning this is typically computed per contig per sample.

**Single-coverage**: Binning using one coverage profile (one metagenome), meaning one coverage vector per contig.

**Multi-coverage**: Binning using multiple coverage profiles for the same contigs across multiple related metagenomes, producing a multi-sample coverage vector per contig.

**Ground truth**: In synthetic datasets, the known origin genome for each read and contig.

**Contig-origin mapping**: Assigning each contig to its source genome (in synthetic datasets) and using those labels to compute bin purity and recovery metrics.

**Clean Bin Bases**: Fraction of binned bases that originate from the dominant source genome for that bin, computed from contig-origin labels.

**Genome representation**: Fraction of ground-truth genomes that are recovered above a defined minimum (define the criterion you used, for example any bin containing >X% of a genome).

**Sequence recovery**: Fraction of total ground-truth bases recovered into bins, computed from contig-origin labels.

**CheckM2**: A machine-learning based estimator of microbial genome completeness and contamination using marker gene features.

**EukCC**: A eukaryote-focused genome quality estimator that reports completeness (and, depending on mode, additional QC signals) using eukaryotic marker sets.

**GTDB**: Genome Taxonomy Database. A genome-based taxonomy for bacteria and archaea.

**GTDB-Tk**: Toolkit that assigns GTDB taxonomy to genomes based on marker genes and phylogenomic placement.

### 5.8. Use of AI tools disclosure

The authors acknowledge the use of AI assistants for both code development and manuscript preparation. Algorithmic and documentation contributions (as detailed in the Methods section) were provided by Anthropic’s Claude-Sonnet. Structural feedback on the manuscript was additionally provided by Google’s Gemini 3 pro. All AI-assisted work was directed, reviewed, and validated by the human authors.

## 6. Data availability

Details on the real metagenome samples used in this project can be found in Supplementary Table S4. Metagenome files corresponding to the samples used in the benchmarking can be downloaded from the JGI Data Portal at https://data.jgi.doe.gov/. Metagenome files corresponding to the samples used in the ground-truth data benchmarking can be downloaded from Zenode https://zenodo.org/communities/quickbin-data. Reference databases included the `GTDB-Tk` data package for GTDB release r226, the `EukCC` database v1.2, the `CheckM2` database, all are publicly available.

## 7. Code availability

`QuickBin` is freely distributed as part of the `BBTools` suite. Source code and installation instructions are available through the `BBTools` website (https://bbmap.org). This includes links to the GitHub repository, SourceForge image, or prebuilt BBTools Docker image available on Docker Hub, enabling reproducible execution in local, cluster, and cloud environments. `BBTools` is free for unlimited use under a LBNL open source license.

Details on version, installation method, and running methods for software and databases used in this project can be found in Supplementary Table S3. In brief, software used in this study and corresponding versions were as follows: Binning was performed with `QuickBin` (BBTools 39.60; deployed via `docker` image `bryce911/bbtools:39.60`), `MetaBAT2` v2.18, `SemiBin2` v2.2.0, `COMEBin` v1.0.4, and `VAMB` v5.0.4; all binners and their respective dependencies, including `CUDA` libraries required for GPU usage, were installed via `pixi` unless otherwise noted. Quality assessment and filtering used `CheckM2` v1.1.0 (installed via `pixi`) and `EukCC` v2.1.3 (executed from the container quay.io/microbiome-informatics/eukcc:2.1.3), and `gradebins` from `BBTools` (v39.54; deployed via the `docker` image at `bryce911/bbtools:39.54`). Taxonomic assignment used `GTDB-Tk` v2.4.1 (installed via `mamba` environment). Read mapping processing used `samtools` v1.22.1 (installed via `pixi`).

## 8. Acknowledgements

The work conducted by the U.S. Department of Energy Joint Genome Institute (https://ror.org/04xm1d337), a DOE Office of Science User Facility, is supported by the Office of Science of the U.S. Department of Energy operated under Contract No. DE-AC02-05CH11231. This research used resources of the National Energy Research Scientific Computing Center (NERSC) (https://ror.org/05v3mvq14), a U.S. Department of Energy Office of Science User Facility located at Lawrence Berkeley National Laboratory.

## 9. Author contributions

BB and JCV designed the overall project. BB developed `QuickBin`. BB generated the synthetic metagenome data, and trained Neural Network models. JCV collected and processed metagenome data for the real metagenome data. JCV installed and troubleshooted software to be run on CPU and GPU nodes at NERSC Perlmutter. JCV wrote the binning pipeline for all binners and performed benchmarking on all data. JCV drafted and wrote the manuscript. BB wrote the methods describing the `QuickBin` algorithm and the generation of synthetic data. All authors edited and accepted the manuscript before submission.

## 10. Competing interests

All other authors declare no competing interests.

## 11. Additional information

The online version contains supplementary material.

## Notes

https://bbmap.org/tools/quickbin

### Competing Interest Statement

The authors have declared no competing interest.

### Summary of Updates

This version of the manuscript has been revised to update the Figure 3 and Figure 4 results.

